# RPTPγ is a redox-regulated suppressor of promigratory EGFR signaling

**DOI:** 10.1101/2022.06.01.494340

**Authors:** Maitreyi S. Joshi, Angel Stanoev, Birga Soetje, Jan Huebinger, Veronika Zorina, Lisaweta Roßmannek, Kirsten Michel, Philippe I. H. Bastiaens

## Abstract

Spatially-organized interaction dynamics between proto-oncogenic epidermal growth factor receptor (EGFR) and protein tyrosine phosphatases (PTPs) determine EGFR’s phosphorylation response to growth factors and thereby cellular behavior within developing tissues. We show here, that and how the coupling between EGFR and RPTPγ activity leads to migratory signaling responses to very low, physiological growth factor stimuli while suppressing aberrant, spontaneous signaling. Single cell imaging of EGFR phosphorylation and PTP oxidation revealed that RPTPγ fully suppresses spontaneous EGFR phosphorylation, while EGF-induced NADPH-oxidase activity enables promigratory signaling responses at the plasma membrane by H_2_O_2_-mediated oxidative inhibition of RPTPγ’s phosphatase activity. The EGF-dependent toggle switch dynamics between interacting EGFR monomers and RPTPγ thereby enables autocatalytically amplified phosphorylation responses to very low, physiological, EGF levels even at sparse receptor expression. This signaling mechanism is distinct from the proliferative signaling stemming from liganded endosomal EGFR complexes at high growth factor concentrations. Accordingly, RPTPγ knock-out results in spontaneous promigratory EGFR signaling but loss of proliferative signaling. We thereby provide evidence of RPTPγ’s suppressor function of oncogenic, promigratory EGFR-signaling from the plasma membrane.

## Introduction

The receptor tyrosine kinase EGFR (Cohen *et al*, 1980; Yarden & Schlessinger, 1987) is implicated in embryonic development, tissue homeostasis and wound healing (Yu *et al*, 2010; Sibilia *et al*, 2007) while EGFR overexpression and hyper-activation through genetic alterations have been linked to malignant transformation (Rowinsky, 2004). Evidence has accumulated that EGFR is embedded in a spatial-temporally organized protein tyrosine phosphatase (PTP) network in order to sense and differentially respond to evolving growth factors patterns in collective cell responses like tissue regeneration (Brüggemann *et al*, 2021; Stallaert *et al*, 2018; Koseska & Bastiaens, 2020; Hiratsuka *et al*, 2015; Lin *et al*, 2022; Hino *et al*, 2020). During this process, proliferative, differentiating or migratory cell responses to ligands of EGFR are spatially-temporally coordinated by the evolving tissue itself that changes the communication between cells. EGFR thus operates within a growth factor sensory system that detects time-varying growth factor patterns by ligand-induced conversion of EGFR monomers to transient dimers that catalytically and autocatalytically generate an amplified phosphorylation response in EGFR monomers (Koseska & Bastiaens, 2020; Baumdick *et al*, 2018; Stanoev *et al*, 2018; Verveer *et al*, 2000; Reynolds *et al*, 2003). These ligandless (auto)-catalytically activated monomers are dynamically maintained at the PM by recycling through the recycling endosomes (REs) in contrast to stabilized liganded ubiquitinated complexes of EGFR that uni-directionally traffic from early endosomes (EEs) to late endosomes (LEs) to be degraded in lysosomes (Baumdick *et al*, 2015; Stanoev *et al*, 2018; Koseska & Bastiaens, 2020). Because of this differential trafficking of liganded ubiquitinated EGFR complexes and (auto)-catalytically activated ligandless monomers, evolving growth factor patterns can generate contextual cellular responses by altering the dynamically established spatial distribution of EGFR between the plasma membrane (PM) and endosomal compartments (Brüggemann *et al*, 2021; Stallaert *et al*, 2018). At the PM, (auto)-catalytic phosphorylation cascades can fully activate the receptor in the absence of growth factors, requiring protein tyrosine phosphatases (PTPs) to counter spontaneous phosphorylation of EGFR (Tischer & Bastiaens, 2003; Koseska & Bastiaens, 2020). In this respect, reciprocal genetic perturbations combined with *in situ* EGFR phosphorylation imaging has identified PM-localized RPTPγ and endoplasmic reticulum associated TCPTP as major dephosphorylating activities that regulate the phosphorylation response of EGFR (Stanoev *et al*, 2018; Koseska & Bastiaens, 2020). However, the 2-3 orders of magnitude higher catalytic activity of PTPs with respect to the kinase activity of EGFR (Fischer *et al*, 1991; Lammers *et al*, 1993) suppresses growth factor responses, requiring EGFR-signaling-induced PTP inhibition to facilitate an EGFR phosphorylation response to growth factors (Denu & Tanner, 1998; Reynolds *et al*, 2003; Bae *et al*, 1997; Stanoev *et al*, 2018).

Herein, we solve this conundrum by showing that EGFR-coupled NADPH-oxidase (NOX1-3) activity enables an amplified EGFR phosphorylation response to low EGF levels by H_2_O_2_-mediated oxidation of the catalytic cysteine of RPTPγ. The resulting switch-like response at the PM is enabled even at low EGFR expression levels due to the interaction between co-recycling RPTPγ and EGFR monomers. Vesicular recycling of RPTPγ through the thiol-reducing environment of the cytoplasm thereby reverses RPTPγ oxidation, closing the redox cycle in the coupled EGFR-RPTPγ toggle switch system at the PM. On the other hand, the constitutive dephosphorylating activity of the PM-proximal pool of ER-associated TCPTP not only safeguards against spurious activation of the ROS coupled EGFR-RPTPγ growth factor sensing system, but also poises EGFR-RPTPγ toggle switch dynamics in a reversible growth factor response regime. Consistent with these findings, knock-out of RPTPγ leads to a highly morphing migratory phenotype of MCF7 cells whereas knock-out of the NOX1-3 subunit p22^phox^ results in a hyperproliferative and a more stationary cellular behavior. RPTPγ is thus a suppressor of promigratory oncogenic EGFR signaling from the PM but not of proliferative cytoplasmic signaling of internalized liganded receptor complexes.

## Results

### EGFR phosphorylation response is dependent on RPTPγ and NOX activity

In order to reveal how the two major EGFR tyrosine phosphatases affect EGFR phosphorylation response, single cell dose-response profiles were obtained after reciprocal genetic perturbation of RPTPγ (*PTPRG*) or TCPTP (*PTPN2*). This was performed by CRISPR-Cas9 knock out (Fig EV1A and EV1B), or by ectopic over-expression of PTP fusion construct with mTFP in corresponding KO-cells. Single cell EGF-dose/EGFR phosphorylation response profiles were obtained in breast cancer-derived MCF7 cells by mapping the fraction of phosphorylated (pY1086, pY1148) EGFR-mCitrine (α_p_) that interacts with PTB-mCherry (phosphotyrosine-binding domain fused to mCherry) (Offterdinger *et al*, 2004) by global analysis of the spatial distribution of fluorescence decay profiles obtained by TCSPC-FLIM (Fig 1A) (Verveer *et al*, 2000; Grecco *et al*, 2009). These low EGFR expressing cells (~10^3^/cell) were co-transfected with PTB-mCherry and EGFR-mCitrine, elevating EGFR-expression to the level of the related MCF10A cells (~1.9±1.7×10^5^ receptors/cell, Fig EV1D) (Stanoev *et al*, 2018). Step-wise EGF-Alexa647 increments enabled imaging receptor occupancy with EGF (α_L_) as function of EGF dose by normalizing the ratiometric fluorescence of Alexa647/mCitrine to that at saturating EGF-Alexa647 dose (Reynolds *et al*, 2003).

**Figure 1.**
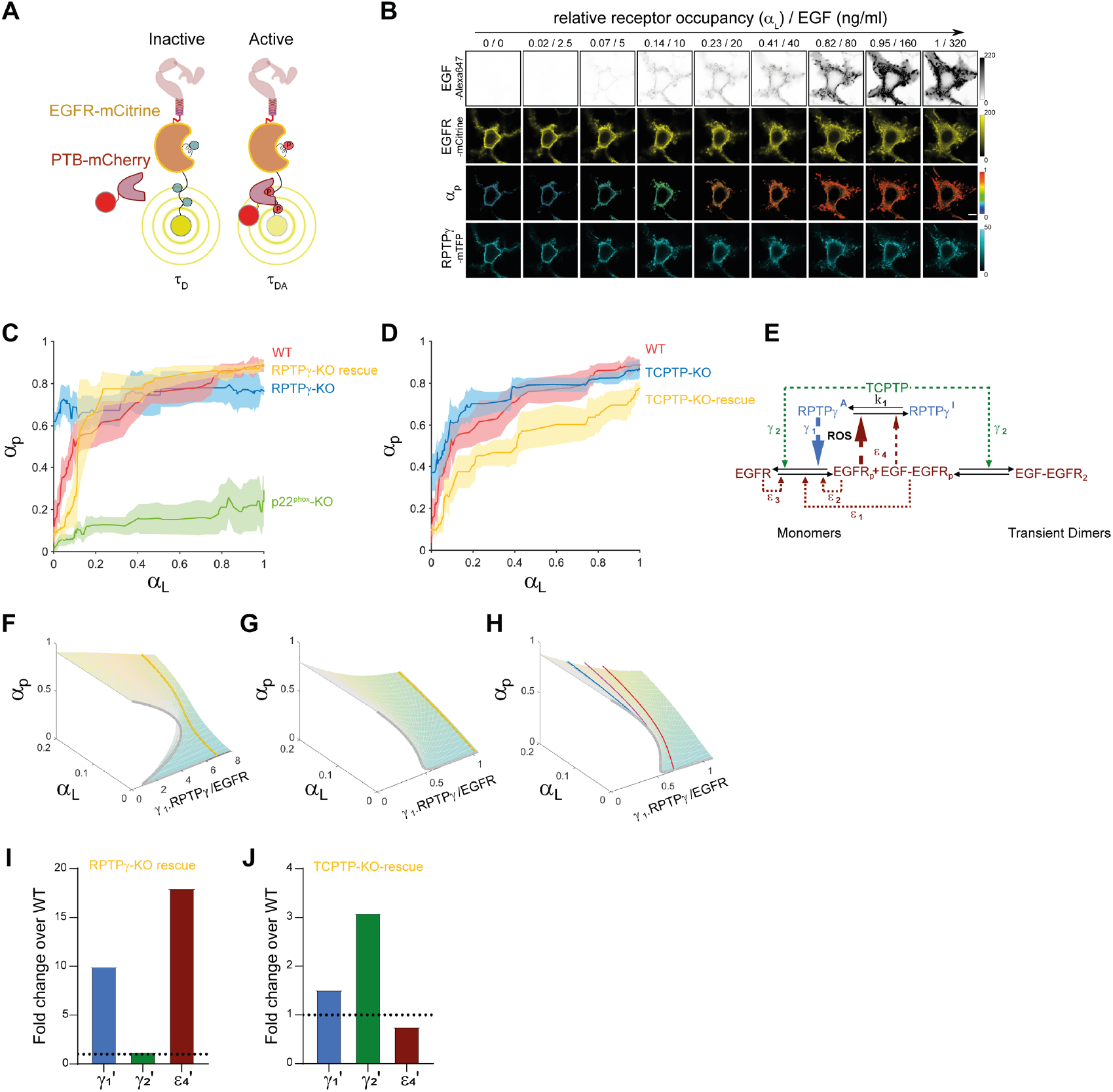
*In cell* dose-response analysis reveals toggle switch dynamics for EGFR phosphorylation. A. Quantitative imaging of EGFR phosphorylation: Binding of PTB-mCherry (acceptor) to phosphorylated EGFR-mCitrine (donor) induces FRET between donor and acceptor resulting in a reduced excited state lifetime (τ_DA_) of the EGFR-mCitrine/ PTB-mCherry complex. Unphosphorylated EGFR-mCitrine exhibits a discrete fluorescence lifetime (τ_D_) that is distinct from τ_DA_. The spatially invariant τ_DA_ and τ_D_ are shared global parameters that enable the mapping of the fraction of phosphorylated EGFR-mCitrine (α_p_, local parameter) within living cells by global analysis. B. Representative fluorescence micrographs of *in cell* dose-response imaging for EGFR phosphorylation after indicated increments of EGF-Alexa647 (0–320 ng/mL) at 1.5’ interval. First row: EGF-Alex647; Second row: EGFR-mCitrine; Third row: phosphorylated EGFR fraction (α_p_); Fourth row: RPTPγ-mTFP; Scale bar: 10 μm. C. EGFR-mCitrine phosphorylation (α_p_) plotted as a function of EGF-receptor occupancy (α_L_) at the PM to incremental EGF-Alexa647 doses in WT (red), RPTPγ-KO (blue), RPTPγ-KO with RPTPγ-mTFP ectopic expression (yellow) and p22^phox^-KO (green) MCF7cells D. same as (C) for WT (red), TCPTP-KO (blue) and TCPTP-KO with TCPTP-mTFP ectopic expression (yellow) MCF7cells. Solid lines: moving medians from single cell profiles; shaded bounds: median absolute deviations. N=3-4, n=12-14 E. EGFR-PTP network architecture depicting the state-transitions and regulatory interactions. Solid arrows: molecular state transitions (p: phosphorylation, I: inactive, A: active), dashed arrows: causal links. Kinetic constants, *γ*_1_,*γ*_2_: phosphatase rate constant of RPTPγ and TCPTP, *k*_1_: rate constant for PTP reactivation. *ε*_1_, *ε*_2_ and *ε*_3_: (auto)-catalytic kinase rate constants for EGFR, *ε*_4_: rate constant for EGFR-dependent phosphatase inactivation. EGFR: EGFR monomer; EGFRp: phosphorylated EGFR monomer; EGF-EGFRp: liganded phosphorylated EGFR monomer, EGF-EGFR2: liganded EGFR dimer at 1:2 stoichiometry. F. Experimentally reconstructed 3D-bifurcation diagram showing the dependence of EGFR phosphorylation (α_p_) on γ_1_.RPTPγ/EGFR and EGF-receptor occupancy (α_L_). Bifurcation surface for RPTPγ-KO cells in which RPTPγ-mTFP was ectopically expressed together with EGFR-mCitrine. Yellow line: experimentally derived dose response trajectory. G. Same as (F) for TCPTP-KO cells in which TCPTP-mTFP was ectopically expressed together with EGFR-mCitrine. H. Same as (F) for RPTPγ-KO cells ectopically expressing EGFR-mCitrine. Red line: experimentally derived dose response trajectory of WT MCF7 cells ectopically expressing EGFR-mCitrine. Purple line: experimentally derived dose response trajectory of TCPTP-KO cells ectopically expressing EGFR-mCitrine. Blue line: experimentally derived dose response trajectory of RPTPγ-KO cells ectopically expressing EGFR-mCitrine. I. Fold changes in kinetic parameters 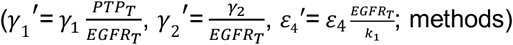 relative to WT for (F). J. Fold changes in kinetic parameters relative to WT for (G).

These α_L_-α_p_ dose-response experiments revealed that EGFR-mCitrine was almost fully phosphorylated in the absence of stimulus in RPTPγ-KO cells (Fig 1C and EV1G). This ‘on-state’ of EGFR was in stark contrast to the ‘off-state’ in unstimulated RPTPγ-KO cells when RPTPγ activity was restored by ectopic expression of RPTPγ-mTFP (Fig 1B and C). In these cells, a clearer sigmoidal switch-like phosphorylation response was attained as compared to WT MCF7 cells, with a defined threshold at ~20% EGF-receptor occupancy (Fig 1C and EV1F). Ectopic overexpression of RPTPγ-mTFP thus accentuates its activity on EGFR against the background of other PTPs. Knock-out of the ER-associated TCPTP resulted in a clear increase in basal EGFR-mCitrine phosphorylation while maintaining a steep response (Fig 1D and EV1I). Ectopic expression of TCPTP-mTFP rescued the low basal EGFR phosphorylation in the absence of stimulus but, however, generated a more gradual hyperbolic phosphorylation response to EGF. This indicates that TCPTP has constitutive suppressive dephosphorylation activity on EGFR (Fig 1D and EV1J). In stark contrast, the EGFR phosphorylation response was highly dampened (Fig 1C and EV1H) in MCF7 cells in which the essential p22^phox^ (*CYBA*) subunit (Ambasta *et al*, 2004) of the plasma membrane NOX1-3 complexes was knocked out (Fig EV1C). This, together with the EGFR phosphorylation ‘on state’ upon RPTPγ KO, points at a toggle-switch relationship between RPTPγ and EGFR, whereby EGFR-activity-induced ROS production inhibits the dephosphorylating activity of RPTPγ.

To scrutinize the respective roles of RPTPγ and TCPTP on the overall phosphorylation response of EGFR, we fitted the dose-response data after reciprocal genetic perturbations with an autocatalytic toggle switch model with EGFR-coupled (RPTPγ-), as well as uncoupled (TCPTP-) phosphatase activity (Reynolds *et al*, 2003; Tischer & Bastiaens, 2003; Stanoev *et al*, 2018) (Fig 1E; methods). The (auto)-catalytic kinetic parameters of EGFR phosphorylation (ε_1_-ε_3_) were considered invariant among the different genetic perturbation experiments and therefore shared among the fits in a global analysis (methods). From the obtained kinetic parameters, a 3D-section of an experimentally accessible 4D bifurcation diagram could be reconstructed for the different genetic perturbations as function of two of the system’s bifurcation parameters (Reynolds *et al*, 2003; Stanoev *et al*, 2018): receptor occupancy α_L_ and RPTPγ/EGFR expression ratio. The third system parameter is the level of constitutive phosphatase activity that will change the 3D-section of the 4D-surface upon its genetic perturbation. This enables to uncover the respective roles of RPTPγ and TCPTP on EGFR phosphorylation response dynamics by the genetic perturbation-induced changes in the bifurcation surface as well as the relative change in the derived kinetic parameters.

The reconstructed bifurcation diagrams exhibited the typical S-shaped bistable dynamical signature of the toggle switch (Fig 1F-H) (Reynolds *et al*, 2003; Stanoev *et al*, 2018), which was strongly accentuated for RPTPγ-mTFP rescue (Fig 1F) and became hyperbolic upon TCPTP rescue (Fig 1G). This confirms that the obtained experimental perturbation data can be explained by the depicted model, where RPTPγ forms a toggle switch with EGFR and EGFR-uncoupled, constitutive TCPTP-activity has a modulatory function on the toggle switch dynamics. This was also apparent from the large fold change of the coupled toggle switch kinetic parameters upon rescue of RPTPγ (γ_1_’,ε_4_’; Fig 1I), whereas TCPTP rescue only increased the constitutive phosphatase kinetic parameter (γ_2_’; Fig 1J). The non-redundant role of RPTPγ in the toggle switch with EGFR was further substantiated by the poising of the system in the preactivated state in RPTPγ-KO cells (Fig 1H, compare red and blue lines). However, from the positioning of the system in the monostable reversible regime, away from the irreversible bistable regime in TCPTP-mTFP rescue cells (Fig 1G), it is apparent that constitutive TCPTP-activity ensures that EGFR phosphorylation responses are reversible (Koseska & Bastiaens, 2020; Stanoev *et al*, 2018). The system was however positioned in the preactivated dynamical regime close to the bistable bifurcation in TCPTP-KO cells (Fig EV1L), showing that constitutive TCPTP activity also helps in safeguarding the mutually coupled EGFR-RPTPγ system from self-activating (Fig 1H, purple line).

### The EGF-induced RPTPγ oxidation switch at the plasma membrane

To experimentally assess whether RPTPγ catalytic cysteine oxidation to cysteine sulfenic acid by H_2_O_2_ generates the inhibitory link from EGFR to RPTPγ via NOX activity (Bae *et al*, 1997; Stanoev *et al*, 2018), we developed a live cell assay to measure the fraction of oxidized RPTPγ (α_ox_) as function of receptor EGF-occupancy (α_L_) (Fig 2A and B). For this, the dimedone warhead derived DYn2 (Paulsen *et al*, 2012) was coupled to Atto590-azide by Cu-based click chemistry (Rostovtsev *et al*, 2002) to obtain a cell permeable mCitrine-FRET acceptor probe DyTo that specifically binds to the sulfenic acid form of oxidized cysteines (Fig EV2A). To image RPTPγ catalytic cysteine oxidation, MCF7 cells ectopically expressing RPTPγ-mCitrine and EGFR-mTFP (~1.9±1.7×10^5^ receptors/cell, Fig EV1D) were stimulated with different doses of EGF for 5’, followed by 5’ incubation with DyTo. The spatially resolved fraction of DyTo-bound oxidized RPTPγ-mCitrine (a_ox_) that exhibits FRET from mCitrine to DyTo was then derived from the global analysis of fluorescence decay kinetics of RPTPγ-mCitrine obtained by TCSPC-FLIM (Verveer *et al*, 2000; Grecco *et al*, 2009). Treatment with 8 mM H_2_O_2_ resulted in pronounced RPTPγ-mCitrine oxidation throughout the cells, but not of the catalytically impaired RPTPγ^C1060S^–mCitrine mutant. This shows that Citrine-DyTo FRET imaging indeed detects the reversible oxidation of the catalytic cysteine of RPTPγ (Fig 2C).

**Figure 2.**
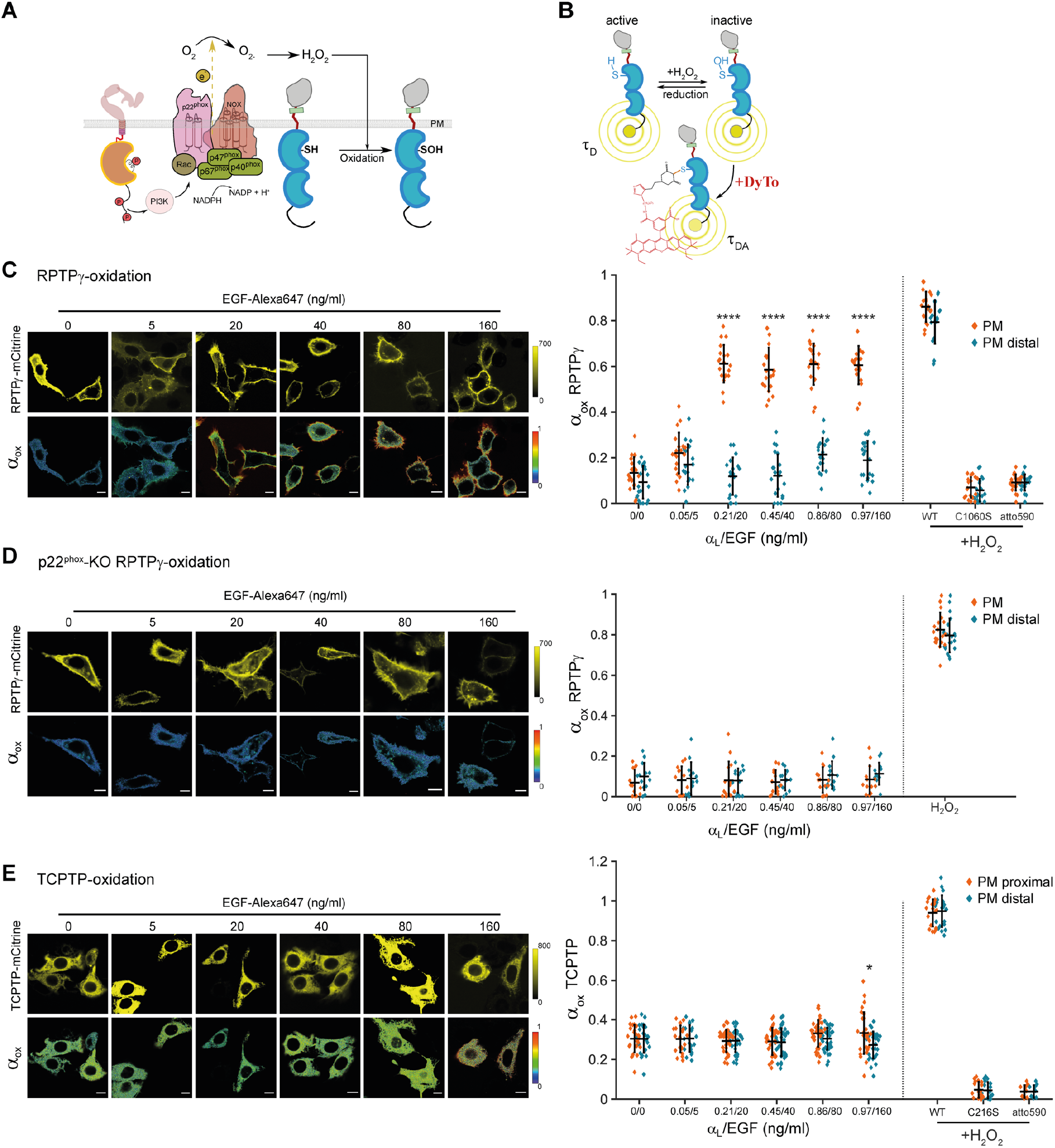
EGF-dependent ROS-generation couples EGFR-phosphorylation to RPTPγ oxidation. A. Reaction schematic of EGFR-dependent PTP-oxidation: Phosphorylated EGFR (red circles) activates PI3K, which results in the activation of Rac-GTPase and the cytosolic components of NOX-assembly like p40^phox^, p47^phox^ and p67^phox^. Recruitment of these components to the PM-based major NOX-unit and p22^phox^ subunit, aids the transfer of electrons from the cytosolic NADPH to extracellular oxygen (O_2_) leading to the formation of superoxide anion (O_2_^-^) that dismutates to hydrogen peroxide (H_2_O_2_). Diffusion of H_2_O_2_ through the PM causes the cysteine oxidation of the PM-vicinal PTPs, from thiol (SH) to sulfenic acid (SOH) state. B. Schematic of FLIM approach for the quantitative imaging of PTP-oxidation in live cells: Binding of DyTo (atto590 acceptor) to oxidized cysteines of PTP-mCitrine (donor) results in FRET between donor and acceptor reducing the excited state lifetime of the donor (τ_DA_). Spatial invariance of τ_DA_ and τ_D_ enable the mapping of the fraction of oxidized PTP-mCitrine (a_ox_, local parameter) by global analysis. C. *in cell* EGF-dose response imaging for RPTPγ-mCitrine oxidation. Left panel: Representativ confocal micrographs of RPTPγ-mCitrine in MCF7 WT cells (top row) together with its oxidized fraction estimated using DyTo-FLIM (a_ox_, bottom row), upon 10’ stimulation with EGF-Alexa647 (0-160 ng/ml). Scale bar: 10 μm. Right panel: Quantification depicting the PM-proximal (orange) and PM-distal (blue) oxidized fractions as functions of receptor occupancy α_L_) and corresponding EGF-Alexa647, or H_2_O_2_ concentration in cells co-expressing EGFR-mTFP, RPTPγ-mCitrine or RPTPγ^C1060S^-mCitrine. Individual cells with mean±SD, N=3, n=13-15. ****p<0.0001: unpaired two-tailed t-test, between PM-proximal and PM-distal fractions. D. Same as in (C), for RPTPγ-mCitrine oxidation in p22^phox^-KO cells. N=3, n=14-26 cells per condition. E. Same as in (C), for TCPTP-mCitrine or TCPTP^C216S^-mCitrine oxidation in MCF7 WT cells. N=3, n=18-21; *p<0.05: unpaired two-tailed t-test, between PM-proximal and PM-distal fractions.

Imaging RPTPγ-mCitrine catalytic cysteine oxidation as function of EGF dose (5-160 ng/ml) in MCF7 cells that co-express EGFR-mTFP (~1.9±1.7×10^5^ receptors/cell, Fig EV1D), showed that a similar toggle-switch response as observed for EGFR phosphorylation occurred for RPTPγ catalytic cysteine oxidation (Fig 1C). Here, RPTPγ-mCitrine oxidation switched at ~20% EGF-receptor occupancy (20 ng/ml EGF-Alexa647) from ~20% to ~60% (Fig 2C, orange symbols). This EGFR-activity-coupled catalytic impairment of RPTPγ was localized to the PM, whereas RPTPγ was in its enzymatically active, reduced catalytic cysteine state in endocytic vesicles (Fig 2C, blue symbols). This indicates that EGF-induced H_2_O_2_ production and the resulting catalytic cysteine oxidation to sulfenic acid was highly confined to the PM proximal area. Consistently, in p22^phox^-KO cells, no RPTPγ-mCitrine oxidation could be detected in response to increasing EGF doses, whereas 8 mM H_2_O_2_ treatment resulted in full RPTPγ-mCitrine oxidation throughout these cells (Fig 2D).

We similarly assessed catalytic cysteine oxidation of ER-associated TCPTP in MCF7 cells that ectopically express TCPTP-mCitrine/EGFR-mTFP (Fig 2E). Imaging TCPTP oxidation as function of EGF dose (5-160 ng/ml) showed a basal and spatially even oxidation independent of EGF stimulus. Only at the highest doses of EGF, a slight but significant oxidation increase in a PM-proximal pool of ER-associated TCPTP could be detected. This confirms that the EGF-induced H_2_O_2_ gradient that is produced by plasma membrane associated NOX1-3 activity is confined to the immediate vicinity of the PM, likely due to the highly reducing environment of the cytoplasm that contains antioxidant systems like peroxiredoxin-thioredoxin (Rhee, 1999; Rhee *et al*, 2005; Woo *et al*, 2010). On the other hand, the overall low but significant level of TCPTP catalytic cysteine oxidation throughout the ER was likely caused by constitutive ER-associated NOX activity (Nisimoto *et al*, 2010). These results show that TCPTP maintains its constitutive dephosphorylating activity during growth factor stimulation, exhibiting a low level of oxidation that is largely uncoupled from EGF-induced EGFR activity. The EGF-induced steep ROS gradient that emanates from the PM thereby constrains catalytic cysteine oxidation to RPTPγ.

### EGFR-RPTPγ interaction conserves phosphorylation response dynamics at low EGFR expression

In order to investigate whether switch-like EGFR activation with low EGF threshold is also maintained at the lower end of EGFR expression range, we measured endogenous EGFR phosphorylation response and resulting down-stream Akt and Erk signaling in MCF7 cells (~10^3^ receptors). We found a switch-like phosphorylation response (pY1068) that occurs at very low receptor occupancy (~1.3 %). This low threshold (<1 ng/ml EGF) amplified response was also apparent in the early (5’) down-stream EGF-dependent activation of Akt and Erk (Fig 3A, middle and right panels). This shows that the γ_1_.RPTPγ/EGFR bifurcation parameter setting in these cells with low EGFR expression poises the system close to the bistable regime (as for ectopic rescue of RPTPγ, Fig 1F). By this, the cells are able to maintain a non-active state below a threshold and generate a highly amplified phosphorylation response to very low, above-threshold level growth factor stimuli.

**Figure 3.**
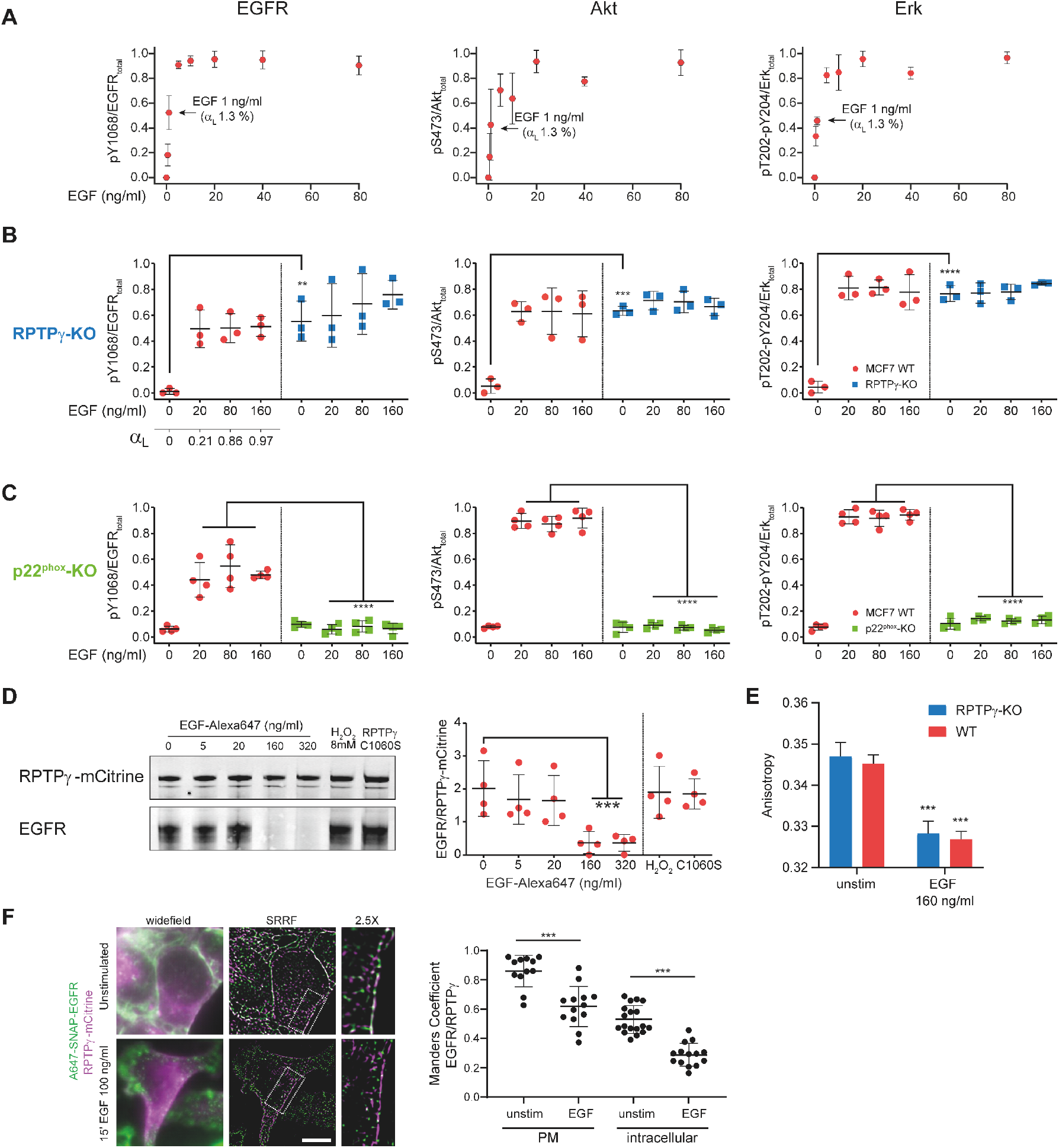
RPTPγ-EGFR monomer interaction enables switch-like responses at low EGFR expression. A. **EGFR (**pY1068)**, Erk (**pT202 and pY204) **and Akt (**pS473) phosphorylation response in WT MCF7 cells as function of EGF concentration derived from Western blot analysis. mean±SD, N=2. B. Same as (A) comparing WT (red), to RPTPγ-KO (blue) MCF7 cells. Normalized phosphorylation versus EGF dose (ng/ml) together with corresponding EGF-receptor occupancy (α_L_) is shown without, and upon 5’ stimulation with different doses of EGF-Alexa647. mean±SD, N=3. **p<0.01, ***p<0.001, ****p<0.0001: unpaired two-tailed t-test. C. Same as (B) comparing WT (red) to p22^phox^-KO (green) MCF7 cells. mean±SD, N=4. ****p<0.0001: unpaired two-tailed t-test. D. Representative IP-western blot and quantification showing co-IP of EGFR (lower blot) upon RPTPγ-mCitrine (upper blot: lanes 1-6) or RPTPγ^C1060S^-mCitrine (lane 7) pull-down by anti-GFP antibody from MCF7 cell lysates: without stimulus (0 ng/ml), upon 10’ stimulus with EGF-Alexa647 (5-320 ng/ml) or 8mM of H_2_O_2_. mean±SD, N=4, ***P<0.001: unpaired two-tailed t-test. E. Quantification of live cell fluorescence anisotropy microscopy measurements of EGFR-QG-mCitrine dimerization state in WT (red) and RPTPγ-KO (blue) MCF7 cells before and after 160 ng/ml EGF-Alexa647 stimulus for 15’. mean ± SEM, N=3, n=31 cells. ***p<0.001: paired two-tailed t-test, against respective unstimulated cases. F. Left panel: Dual-color widefield images (first column), SRRF reconstructions (second column) with magnifications of boxed areas (third column) of Alexa647-SNAP-EGFR (green)/RPTPγ-mCitrine (magenta) of cryo-arrested MCF7 cells, unstimulated (top row) or stimulated with 100 ng/ml EGF (bottom row) for 15’. Scale bar: 10 μm. Right panel: corresponding Manders colocalization coefficients for Alexa647-Snap-EGFR/RPTPγ-mCitrine from SRRF reconstructions on intracellular compartments or PM area for unstimulated (n=12-18) and 15’ EGF-stimulated (n=13-14) cells. ***P<0.001: unpaired two-tailed t-test.

In these low EGFR expressing cells, knock-out of RPTPγ also led to high endogenous EGFR, Akt and Erk phosphorylation levels (Fig 3B and EV2B) whereas p22^phox^-KO cells maintained the EGFR phosphorylation off-state, irrespective of growth factor stimulus (Fig 3C and EV2C). These results support the central role of the ROS-mediated EGFR-RPTPγ toggle switch in phosphorylation responses to low growth factor levels even at low receptor expression levels. Furthermore, they show that ligandless, autocatalytically amplified, phosphorylated EGFR-monomers can engage down-stream Erk and Akt signaling from the PM. Also, in a colorectal cancer cell line (HT29), known to be deficient in RPTPγ expression due to promoter methylation (Van Niekerk & Poels, 1999), a high basal EGFR phosphorylation could be significantly lowered by ectopic expression of RPTPγ (Fig EV2D-F). This indicates that RPTPγ is a potential tumor suppressor of aberrant EGFR signaling.

To examine the possibility that a switch-like response to low level EGF stimuli is facilitated by enhancing the catalytic efficiency of RPTPγ activity on EGFR by direct interaction, we carried out immunoprecipitation of RPTPγ-mCitrine and measured the extent of EGFR pull-down as function of EGF dose (Fig 3D and EV2G). An interaction between EGFR and RPTPγ-mCitrine was indeed apparent in the absence of stimulus as well as at low, sub-saturating doses of EGF (5 and 20 ng/ml), conditions where (auto)-catalytically phosphorylated monomers predominate the EGFR response. This interaction did however not occur upon treatment with receptor-saturating EGF doses (160 and 320 ng/ml) that predominantly generates stabilized liganded EGFR complexes. We verified by homo-FRET measurements using fluorescence anisotropy microscopy on an EGFR-QG-mCitrine construct (Baumdick *et al*, 2015), that saturating EGF indeed generates EGF-EGFR complexes in RPTPγ-KO cells similar to that in WT cells and corroborated that (auto)-catalytically activated, phosphorylated EGFR in RPTPγ-KO cells is predominantly monomeric (Fig 3E). RPTPγ thus constitutively interacts with EGFR monomers but not with stabilized liganded EGFR complexes.

To investigate whether this change in EGFR-RPTPγ interaction dynamics is reflected in the nanoscale organization of EGFR (Ibach *et al*, 2015; Masip *et al*, 2016) in relation to RPTPγ, we assessed the colocalization of Alexa647-SNAP-EGFR and RPTPγ-mCitrine after receptor-saturating EGF stimulation by super resolution radial fluctuation microscopy after ultra-rapid cryo-arrest on a fluorescence microscope (Huebinger *et al*, 2021). In serum-starved, quiescent cells, a co-organization of EGFR nanoclusters within larger RPTPγ patches along the PM could be observed. However, upon 15’ 100 ng/ml EGF stimulus, EGFR clusters segregated from RPTPγ at the PM as well as in endocytic structures (Fig 3F). This change in EGFR-RPTPγ nanoscale organization upon receptor-saturating EGF stimulus confirms that the interaction between unliganded EGFR monomers and RPTPγ at the PM is disrupted upon formation of liganded EGFR complexes.

### Recycling RPTPγ-EGFR generates a growth factor responsive spatial redox cycle

Our results indicated that the loss in the interaction between EGFR monomers and RPTPγ upon high EGF stimulus affects the spatial organization of EGFR in relation to RPTPγ. In order to investigate if this has a function for growth factor sensing, we first measured the intracellular re-distribution of ectopically expressed RPTPγ-mCitrine and EGFR-mCherry by immunostaining against endosomal markers for early-(EEA1), recycling-(Rab11a) and late endosomes (Rab7). Prior to EGF stimulus, RPTPγ-mCitrine and EGFR-mCherry exhibited co-localization on the PM as well as the RE (Fig 4A and B, left top-bottom panels), indicating constitutive co-recycling of both proteins. After saturating EGF stimulus (160 ng/ml), which mostly generates liganded EGFR complexes, RPTPγ still localized to the PM and RE (Fig 4A and B, left and middle panels). However, EGFR separated from RPTPγ, trafficking towards the LE via the EE (Fig 4B, middle and right panels, EV3A and B). This is consistent with the dynamically maintained constitutive co-recycling of RPTPγ-EGFR being disrupted upon the generation of EGF-induced EGFR complexes that unidirectionally traffic from the PM, via EEs to the LE. In order to substantiate that RPTPγ is dynamically maintained at the PM by vesicular recycling, we enhanced the biogenesis of the RE by ectopic expression of BFP-Rab11a. This caused the steady-state distribution of RPTPγ-mCitrine to shift towards the RE in dependence on the level of BFP-Rab11a expression (Fig 4C and D). RPTPγ vesicular recycling was also apparent from the time-dependent re-equilibration to steady-state at the PM of paGFP-RPTPγ photoactivated on the RE (Fig 4E, EV3D and E).

**Figure 4.**
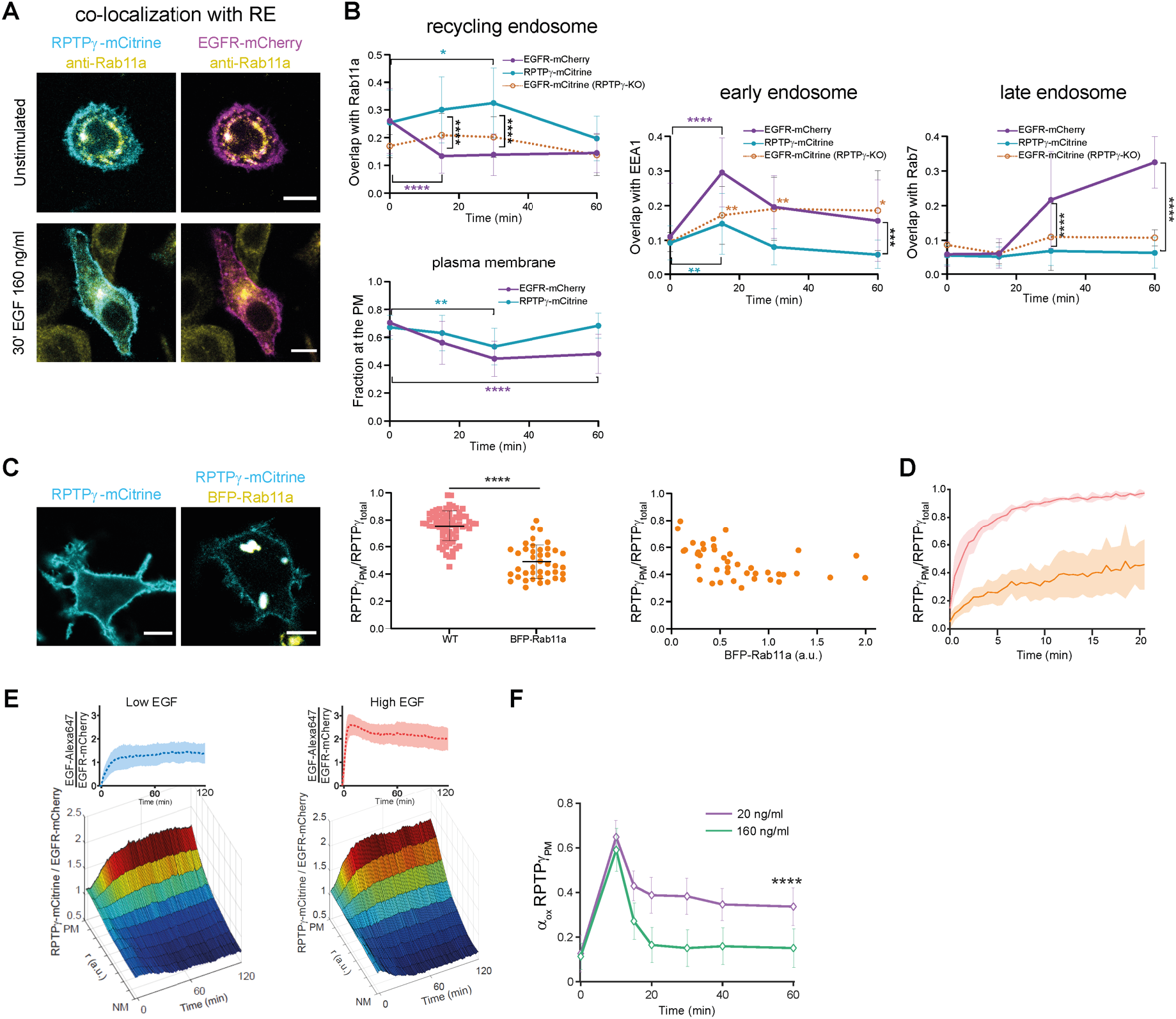
EGFR-RPTPγ redox-phosphorylation-cycles are spatially coupled. A. Representative confocal micrographs of MCF7 WT cells showing the co-localization of RPTPγ-mCitrine (cyan, first column) and EGFR-mCherry (magenta, second column) with recycling-endosome marked by immunostaining against Rab11a (yellow), without- (top row) or after 30’ EGF-DyLight405 (160 ng/ml) stimulus (bottom row). Scale bar: 10 μm. B. Fraction of RPTPγ-mCitrine (cyan) or EGFR-mCherry (magenta) that spatially overlaps with Rab11a-positive recycling endosomes (top left), PM (bottom left), EEA1-positive early endosome (middle) or Rab7-positive late endosome (right) in MCF7 cells as function of time after 160 ng/ml EGF-stimulus. Orange symbols/dotted line: the same for EGFR-mCitrine in RPTPγ-KO cells. For all conditions: N=3, n=25-30, *p<0.05, **p<0.01, ***p<0.001, ****p<0.0001: unpaired two-tailed t-test, colored: versus 0’; black: between RPTPγ or EGFR. C. Representative confocal micrographs (left) and quantification (middle) of MCF7 WT cells depicting the steady state localization of RPTPγ-mCitrine (cyan), without or with co-expression of BFP-Rab11a (yellow). The change in PM-fraction of RPTPγ-mCitrine as a function of BFP-Rab11a expression as measured by fluorescence intensity (right). Scale bar: 10 μm. N=2, n>40; ****P<0.0001: unpaired two-tailed t-test. D. Redistribution of paGFP-RPTPγ to the PM post perinuclear photoactivation, in cells with (orange) or without (pink) the co-expression of BFP-Rab11a. Scale bar: 10 μm. N=3, n=4-7. E. Average spatial-temporal maps constructed from confocal micrographs obtained at 1’ interval from live MCF7 WT cells showing the distributions of RPTPγ-mCitrine/EGFR-mCherry as a function of their radial distance and time, upon sustained low (left panel, 20 ng/ml, N=3, n=13) or high (right panel, 160 ng/ml, N=3, n=14) stimulus with EGF-Alexa647. The top inset depicts the time dependent EGF-Alexa647/EGFR-mCherry (proportional to α_L_) fluorescence intensity at the PM. F. Temporal profiles for the oxidized fraction of RPTPγ-mCitrine (a_ox_) at the PM estimated using DyTo-FLIM, upon receptor sub-saturating (20 ng/ml, magenta) or saturating (160 ng/ml, green) sustained EGF-Alexa647 stimulus, obtained from live MCF7 WT cells expressing EGFR-mTFP and RPTPγ-mCitrine. mean±SD, N=3, n=18-23 cells per time point. ****p<0.0001, from 15’ to 60’: unpaired two-tailed t-test between sub-saturating and saturating response.

In RPTPγ-KO cells, however, EGFR did not reach the LE upon saturating EGF stimulation, and instead kept recycling as apparent from the co-localization of EGFR-mCitrine with EE and the RE. The altered EGFR-trafficking upon RPTPγ-KO thus maintains a substantial fraction of EGFR on the PM, irrespective of EGF stimulus, thereby hindering its efficient lysosomal degradation (Fig 4B, dotted profiles). The deregulated EGFR trafficking in RPTPγ-KO cells was also apparent from substantial colocalization of EGFR with the ER (Fig EV3C) and other endomembrane structures as well as strong enrichment on PM ruffles. This indicates that EGFR trafficking to the PM after its biosynthesis and ER-insertion as well as its vesicular dynamics is deregulated in the absence of RPTPγ.

To next relate RPTPγ-EGFR spatial-temporal dynamics to ROS-mediated toggle-switch reaction dynamics, we obtained the temporal profile of RPTPγ catalytic cysteine oxidation in response to a low (20 ng/ml) or high (160 ng/ml) EGF-Alexa647 stimulus and compared this to high-resolution radial spatial-temporal maps (STMs) (Stanoev *et al*, 2018) of EGFR-mCherry/RPTPγ-mCitrine from 2h CLSM (confocal laser scanning microscopy) time-lapse imaging (Fig 4E, F and EV3F-I). In these experiments, the constant supply of EGF continuously generates a fraction of liganded EGFR complexes from recycling EGFR-RPTPγ that thereby separate from RPTPγ and are depleted from the PM. The EGF-induced depletion of EGFR from recycling EGFR-RPTPγ will therefore change the steady-state of the system to a higher fraction of RPTPγ with respect to EGFR at the PM in dependence on EGF concentration. A high (160 ng/ml) EGF stimulus indeed rapidly depleted the recycling EGFR pool from the PM by unidirectional traffic of EGFR complexes to the LE (Fig 4E, right panel and EV3G). However, a low (20 ng/ml) EGF stimulus sustained a higher steady-state amount of recycling EGFR monomers at the PM that rebind EGF at the PM and thereby can re-activate EGFR autocatalysis (Fig 4E, left panel and EV3F).

For the two EGF-stimuli modes (low, high), RPTPγ attained the maximally oxidized, inactive state at the PM within ~10’, the timescale of maximal EGFR-phosphorylation (Fig 4F and EV3H-J). However, whereas the high sustained EGF-stimulus (160 ng/ml) caused RPTPγ oxidation to fall back to basal levels in ~20’ (Fig 4F, green) due to depletion of EGFR from the PM (Fig EV3G, right panel), the low sustained EGF-stimulus (20 ng/ml) caused the dynamic equilibration to a high steady-state oxidation level (Fig 4F, magenta). These results show that the low EGF-induced autocatalytic phosphorylation of continuously recycling EGFR-monomers maintain oxidation of RPTPγ at the PM by sustaining NOX-activity. This is in contrast to the high EGF stimulus, which rapidly depletes signaling active, phosphorylated EGFR complexes from the PM and thereby consumes the available pool of recycling EGFR. Vesicular recycling of RPTPγ-EGFR thus restores the catalytically active state of RPTPγ, which is consistent with the observation that RPTPγ’s catalytic cysteine is in the reduced state in endosomes (Fig 2C).

### RPTPγ is a suppressor of EGFR promigratory signaling response

In order to relate the ROS-regulated EGFR-RPTPγ molecular growth factor sensing system to cellular phenotype, we compared the morphodynamic growth factor response of MCF7 WT cells to those in which either RPTPγ or p22^phox^ was knocked out. First, the dependence of growth factor induced PM morphodynamics on EGFR expression was assessed by comparing endogenous (~10^3^ receptors/cell) to ectopic EGFR expression (analogous to MCF10A, ~1.9±1.7×10^5^ receptors/cell, Fig 5A and EV1D). For endogenous EGFR expression, EGF-induced PM shape-changes were measured over a short time period of an hour by time-lapse fluorescence microscopy of cells expressing the PM marker BFP-tkRas (Fig 5B and C). For ectopic EGFR-mCitrine expression, Citrine fluorescence was used to identify the peripheral PM (Fig EV4A and 5D). Time-integrated morphological profiles of cells were quantified as the distribution in the ratio of the perimeter of an equiareal circle to the actual perimeter of the cell (P_circle_/P_cell_), where P_circle_/P_cell_ approaches one for a circular shape and tends towards zero in case of enhanced overall curvature generated by protrusions.

**Figure 5.**
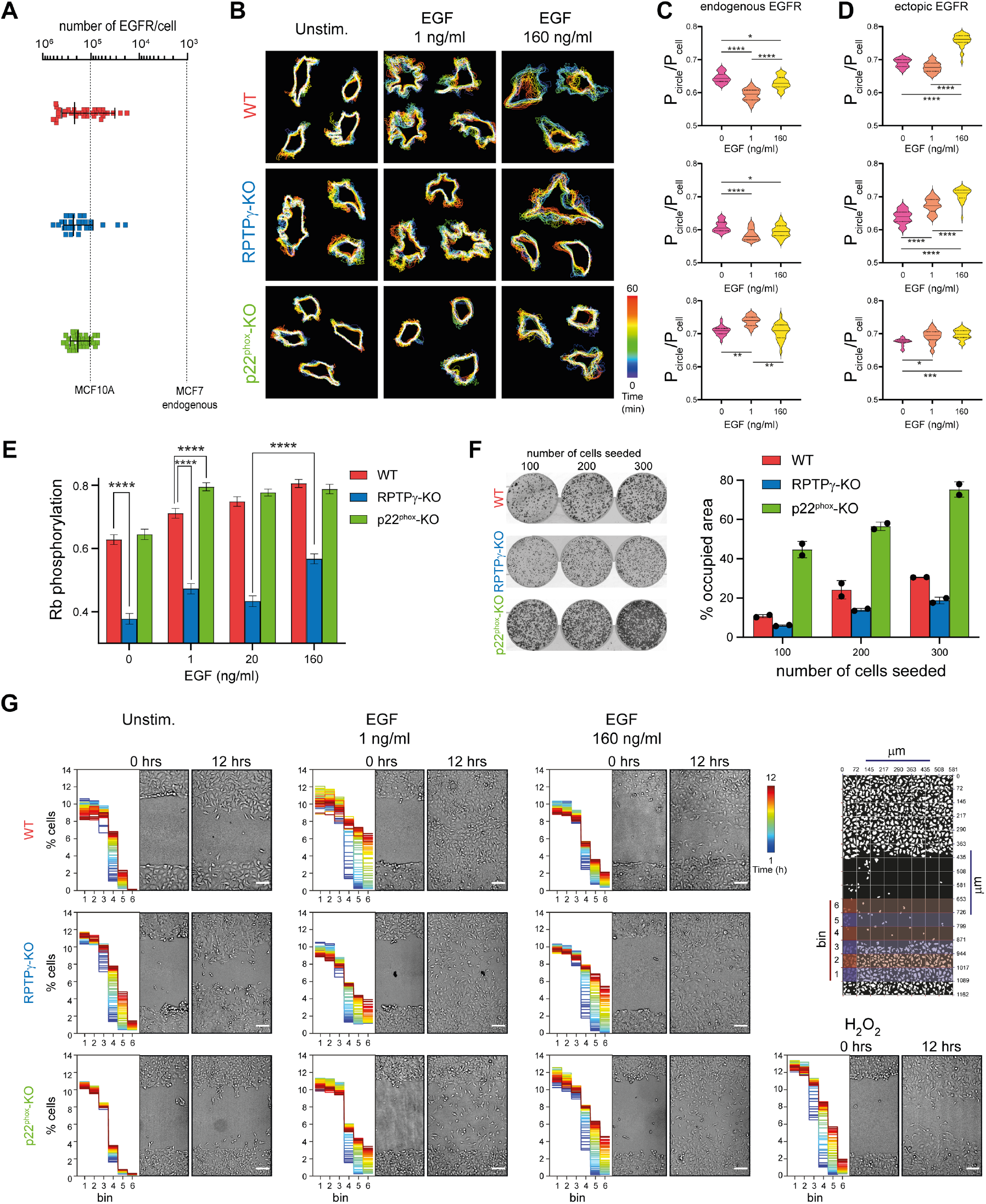
RPTPγ is a suppressor of oncogenic EGFR promigratory signaling. A. Quantification of ectopic EGFR-mCitrine expression in WT (red), RPTPγ-KO (blue) and p22^phox^-KO (green) MCF7 cells. B. Representative cell contour maps showing the temporal changes (color bar: time (min)) in the cell morphology for WT (upper row), RPTPγ-KO (middle row) and p22^phox^-KO (bottom row) MCF7 cells, expressing PM-marker BFP-tkRas without (first column) or during 60’ EGF-Alexa647 stimulus: 1 ng/ml (second column); 160 ng/ml (third column). C. Quantification of EGF-stimulus induced morphodynamics at endogenous EGFR (~10^3^ receptors/cell) in MCF7 cells (n= 9-20), integrated over time by the ratio of the perimeter of an equiareal circle to the actual perimeter of the cells (P_circle_/P_cell_). First row: WT, second row: RPTPγ-KO, third row: p22phox-KO MCF7 cells. D. Same as (C) for MCF7 cells ectopically expressing EGFR-mCitrine (~2×10^5^ receptors/cell). n=11-39. *p<0.05, **p<0.01, ***p<0.001, ****p<0.0001: one-way ANOVA with Šídák’s multiple comparisons E. Quantification of cell proliferation using retinoblastoma (Rb) protein phosphorylation detected by immunofluorescence, for WT (red), RPTPγ-KO (blue) and p22^phox^-KO (green) MCF7 cells without or post 24 h of EGF-Alexa647 treatment (1, 20, 160 ng/ml). mean±SEM., N=2, n>46. ****p<0.0001: unpaired two-tailed t-test. F. Left panel: Clonogenic assay of WT (top), RPTPγ-KO (middle) and p22^phox^-KO (bottom) MCF7 cells plated at an initial density of 100, 200 and 300 cells/well, fixed and imaged on the 7^th^ day post plating. Right panel: mean±SD percentage of culture-well area covered by the cell-colonies. N=2. G. Representative transmitted light micrographs of WT (top row), RPTPγ-KO (middle row) and p22^phox^-KO (bottom row) MCF7 cells, without (first column) and during stimulation with EGF-Alexa 647 (1 ng/ml, second column; 160 ng/ml, third column) or H_2_O_2_ (0.5 mM, fourth column) at the indicated times (0, 12 h) after removal of the migration barrier. Scale bar: 100 μm. Insets on the left: Quantification of average cell number (% over total) over time (color code upper right bar) across 6 spatial bins around the initial cell front (location of the lateral bins in the migration chamber depicted on the right). N=4-5.

Comparison of the PM morphodynamics in the absence of EGF stimuli showed that RPTPγ-KO cells exhibited enhanced and dynamic protrusions with respect to WT cells, whereas p22^phox^-KO cells had less membrane protrusions with strongly reduced dynamics (Fig 5B). This was also apparent from the lower median P_circle_/P_cell_ and wider distribution upon RPTPγ-KO as opposed to a higher median and narrower distribution in p22^phox^-KO relative to WT cells (Fig 5C). The relative morphodynamic changes were similar in MCF7 cells ectopically expressing EGFR-mCitrine (Fig 5C and EV4A). These results are consistent with RPTPγ activity suppressing autonomous promigratory EGFR signaling from the plasma membrane. Both in WT and EGFR-mCitrine expressing cells, low subsaturating (1 ng/ml) EGF-stimulus caused the cells to switch to a highly dynamic morphing behavior that was similar to that of the unstimulated and stimulated RPTPγ-KO cells, whereas p22^phox^-KO cells tended to retract and round off. This is consistent with EGFR-activity inducing NOX1-3 mediated ROS production that is necessary to switch-on promigratory, (auto)-catalytically activated EGFR signaling at the PM by inhibition of RPTPγ. However, high, saturating EGF doses (160 ng/ml), caused WT cells to exhibit retraction and rounding, a phenotypic response that was accentuated in p22^phox^-KO cells. In contrast, RPTPγ-KO cells maintained their membrane morphing dynamics now combined with enhanced retraction.

Since cellular retraction pointed at the initiation of a proliferative response from internalized, liganded receptor complexes (Brüggemann *et al*, 2021), we assessed proliferative signaling for the WT and KO MCF7 cell types by measuring phosphorylation of the S-phase blocker retinoblastoma protein (pRb) by immunofluorescence (Fig 5E). Whereas p22^phox^-KO cells exhibited a hyperproliferative response to growth factors, RPTPγ-KO cells were clearly hypoproliferative, only increasing Rb phosphorylation at saturating EGF stimulus. These findings were independently assessed by clonogenic assays under growth factor serum conditions that provide a measurement of overall proliferation from the area occupied by cell colonies (Fig 5F) (Klein *et al*, 2019). These results are consistent with internalized liganded receptor complexes providing proliferative signals from endosomes whereas autocatalytically activated EGFR at the PM sustains promigratory signaling (Brüggemann *et al*, 2021).

To investigate how the EGFR-RPTPγ system affects long term collective cell behaviour, we investigated tissue regeneration in dependence on exogenous EGF concentration in a simple unicellular MCF7 model system of wound healing (Brüggemann *et al*, 2021). For this, invasion of confluent monolayers of WT as well as the RPTPγ- and p22^phox^-KO cells into a cell-free gap was monitored over 12 h after removal of a barrier without or with (low or high) EGF stimulus (Fig 5G). The evolution of gap closure by the invading cell mass (Fig 5G, left inset and EV4B) was determined by quantifying cell density in lateral bins (Fig 5G, depiction on top right; methods). Full closure could be induced in WT cells upon stimulation with low EGF (1 ng/ml), whereas high EGF (160 ng/ml) inhibited the closure process. In the former case, the right balance between promigratory signaling at the PM and proliferative endocytic signaling, possibly induced by cell-cell contact (Stallaert *et al*, 2018), generates sufficient cell mass that closes the gap in an apparent coordinated way (Movie EV1). In the latter, high EGF case, mostly proliferative signaling from endocytosed receptors led to the loss of a distinct tissue boundary from which individual cells migrated into the gap. In contrast, RPTPγ-KO cells collectively exhibited enhanced migration into the gap already under serum starvation conditions that was only slightly enhanced upon low EGF-stimulus (1 ng/ml). This is consistent with sustained autonomous autocatalytically activated EGFR promigratory signaling from the PM in the absence of the inhibitory RPTPγ phosphatase activity. Due to low proliferating signaling activity in these RPTPγ-KO cells (Fig 5E and F), the diluting migratory cell mass was not able to close the gap in a coordinated way. Altered EGFR distribution in RPTPγ-KO cells (Fig 4B), sustained high membrane activity, which in combination with increased proliferation from internalized liganded receptor complexes at high EGF-dose (160 ng/ml) (Fig 5E) intensified the overall movement of the cell mass that could enter the gap. Although, serum starved p22^phox^-KO cells exhibited only a minimal collective cell-front progression, 0.5 mM H_2_O_2_ induced significant cell-invasion into the gap, phenocopying the collective-migration observed in unstimulated RPTPγ-KO cells. This corroborates that the inhibition of RPTPγ by H_2_O_2_ leads to spontaneous autocatalytic activation of EGFR at the PM and associated promigratory signalling. In contrast, the normally migratory low EGF stimulus caused a minimal collective cell-front progression for p22^phox^-KO cells, where proliferating cells apparently released individual migrating cells into the gap. This collective cell front progression was increased upon high EGF (160 ng/ml) stimulus, likely driven by hyperproliferation.

## Discussion

We provide evidence that the ROS mediated toggle switch coupling of EGFR kinase and the RPTPγ phosphatase is at the core of an (auto)-catalytic growth factor-sensing system that enables promigratory responses at the PM to very low, physiological EGF stimuli. This system operates under the kinetic premise that at low physiological EGF concentrations, transient EGF-EGFR2 dimers (3-10s) (Koseska & Bastiaens, 2020; Ichinose *et al*, 2004; Salazar-Cavazos *et al*, 2020) are reaction intermediates that catalytically and autocatalytically (Baumdick *et al*, 2015; Koseska & Bastiaens, 2020; Stanoev *et al*, 2018; Reynolds *et al*, 2003) generate signaling-competent phosphorylated EGFR monomers (Fig 6, yellow horizontal arrow). Such an (auto)-catalytic system will fully phosphorylate without the opposing dephosphorylating activity of PTPs, which on the other hand, will suppress phosphorylation of EGFR upon growth factor stimulation due to the high catalytic activity of PTPs (Reynolds *et al*, 2003; Tischer & Bastiaens, 2003; Koseska & Bastiaens, 2020). This was resolved by not only establishing that RPTPγ is the major phosphatase that suppresses spontaneous EGFR phosphorylation at the PM, but also that EGFR-activity-coupled ROS production (H_2_O_2_) by NOX1-3 enables EGF-induced phosphorylation responses by inhibition of RPTPγ through oxidation of its catalytic cysteine (Fig 6 black and purple vertical arrows). We could show that the EGFR-RPTPγ phosphorylation-dephosphorylation cycle is dynamically established in space by vesicular recycling of RPTPγ that restores its inhibitory catalytic activity on EGFR by re-reduction of its oxidized catalytic cysteine through the thiol reducing environment of the cytoplasm. This spatial cycle thereby closes the coupled redox- and dephosphorylation-phosphorylation cycles, conveying reversibility to the EGFR-RPTPγ toggle-switch phosphorylation response dynamics (Fig 6).

**Figure 6.**
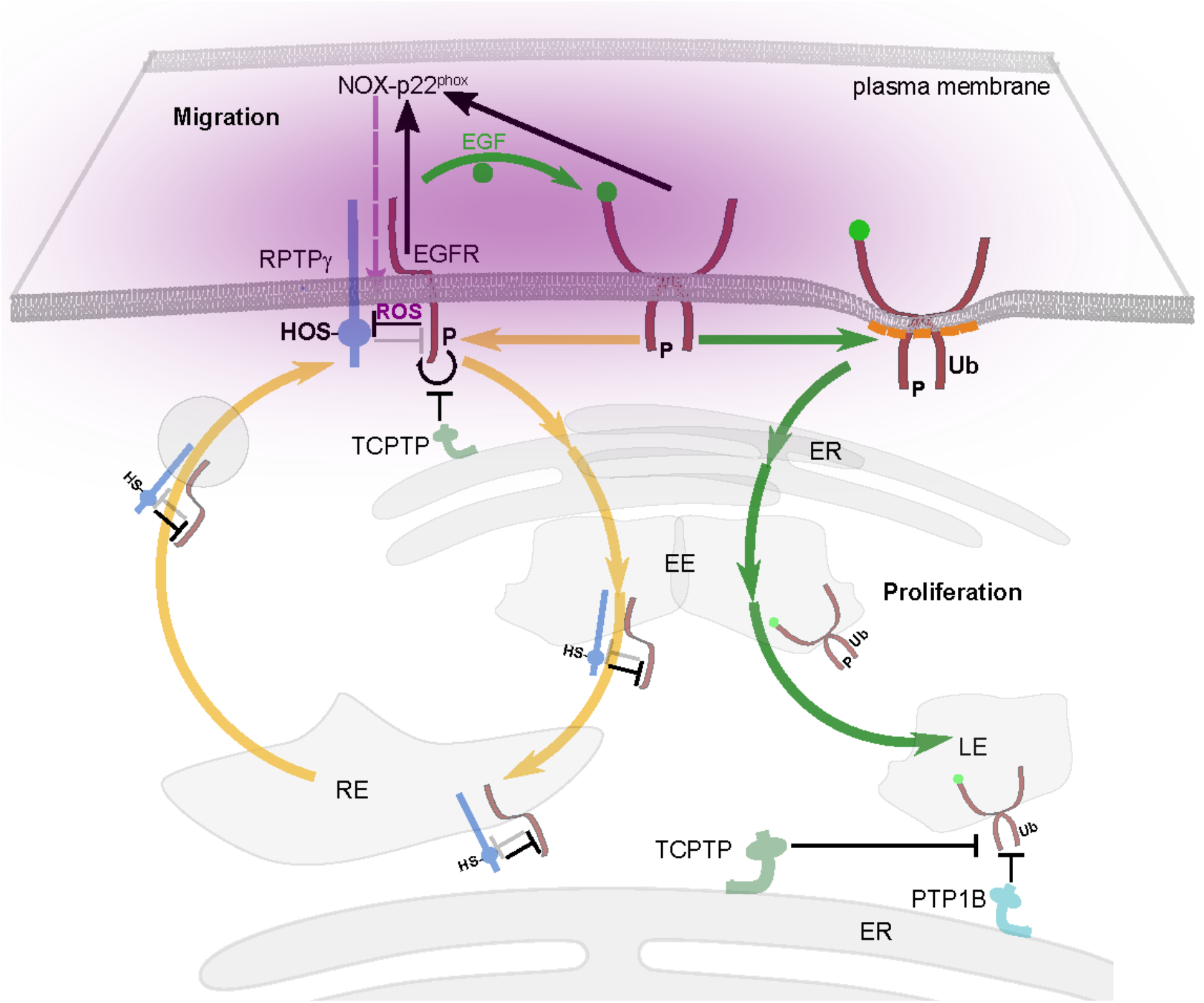
Reaction scheme of the spatially organized EGFR-PTP growth factor sensory system. In the quiescent state the continuous recycling (yellow arrows) of interacting RPTPγ-EGFR monomers maintains reduced (SH) catalytically active state of RPTPγ that dephosphorylates and maintains the catalytically inactive state of EGFR. Upon low level EGF-stimulus (curved green arrow) transient EGF-EGFR2 dimers (auto)-catalytically generate phosphorylated EGFR monomers (yellow straight arrow). These activate NOX complexes (black arrows to NOX-p22^phox^) to locally produce H_2_O_2_ (purple cloud) at the PM that inactivates the inhibitory phosphatase activity of RPTPγ (SOH) on autocatalytic EGFR phosphorylation (black curved arrow), generating promigratory EGFR signaling at the PM. This activation switch is established by the ROS-mediated toggle switch relationship between EGFR and RPTPγ (represented by the mutual inhibition of EGFR and RPTPγ). The inhibitory phosphatase activity of PM-proximal TCPTP maintains reversibility in the response dynamics to EGF together with the recycling through the RE of interacting RPTPγ-EGFR (curved yellow arrows) that reinstates the reduced (SH) catalytically active state of RPTPγ that dephosphorylates EGFR. This ensures that an amplified EGFR phosphorylation response at the PM reverts to basal phosphorylation levels upon depletion of EGF. The liganded transient EGFR dimers can also generate stable, ubiquitinated (Ub) EGFR complexes (green arrow) in dependence on EGF concentration. Higher EGF concentration shifts the balance towards this branch of proliferative EGFR signaling originating from unidirectional trafficking of EGFR complexes through endosomal compartments (green curved arrows). EGFR signal duration is determined by the dephosphorylating activities of ER-associated TCPTP and PTP1B while the receptor complexes traffic to the LE via the EE.

Experimental bifurcation analysis uncovered not only the non-redundant role of RPTPγ in the toggle switch with EGFR, but also that the PM-proximal pool of TCPTP activity poises the system in a reversible ‘ growth factor response regime that additionally safeguards against spontaneous activation of the EGFR-RPTPγ toggle switch. This switch-like EGFR phosphorylation response to low levels of EGF (~1 ng/ml) was conserved at the lower end of EGFR expression range (~10^3^ receptors). The interaction between RPTPγ and EGFR monomers thereby implicates that it is the γ_1_.RPTPγ/EGFR bifurcation parameter (~RPTPγ/EGFR relative expression level at the PM) that determines the steep phosphorylation response to low growth factor levels, positioning the system in, or close to a reversible bistable regime.

This EGFR toggle-switch signaling system at the PM is distinct from the liganded ubiquitinated EGFR I complexes that exhibit endosomal proliferative signaling while they unidirectionally traffic to the LE, via the EE (Brüggemann *et al*, 2021; Stallaert *et al*, 2018) (Fig 6). This branch of internalized EGFR signaling deplete the recycling EGFR pool and is dependent on EGF concentration (Brüggemann *et al*, 2021) as well as EphR mediated cell-cell contact signals that redistribute EGFR to EEs (Stallaert *et al*, 2018). This enables local context dependent proliferative responses in an evolving tissue. Endosomal EGFR signaling also originates from transient liganded EGFR dimers at the PM (Fig 6, green arrows) but is neither dependent on NOX activity nor on interaction with RPTPγ and the temporal signaling response is instead dependent on the ER-associated TCPTP and PTP1B phosphatase activities while receptor complexes unidirectionally traffic to the LE (Haj *et al*, 2002; Yudushkin *et al*, 2007; Stanoev *et al*, 2018). RPTPγ is thus a suppressor of oncogenic migratory EGFR signaling from the PM but not that of proliferative signaling of endocytosed ubiquitinated receptor complexes. Our results provide support that spatial-temporally organized tyrosine-kinase/protein tyrosine phosphatase networks are adaptive growth factor sensing systems that couple higher-scale collective tissue behavior to cellular responses and vice-versa.

## Materials and Methods

### Antibodies

Primary antibodies: rabbit anti-RPTPγ (gift from C. Sorio, Department of Pathology, University of Verona, Verona, Italy), mouse anti-TCPTP (MAB1930, R&D Systems, Minneapolis, MN), mouse anti-p22^phox^ (ab80896, Abcam, Cambridge, UK), anti-cysteine sulfenic acid 2-thiodimedone (ABS30, Merck, Darmstadt, Germany), living colors mouse anti-GFP (632681, Clontech, Mountain View, CA), goat anti-GFP (ab5450, Abcam), rabbit anti-phospho EGFR Y1068 (3777, Cell Signaling Technology, Danvers, MA),, mouse anti-phospho EGFR Y1068 (2236, Cell Signaling Technology) rabbit anti-EGFR (4267, Cell Signaling Technology), goat anti-EGFR (AF231, R&D Systems), rabbit anti-phospho-ERK-1/2 Thr/Tyr 202/204 (9101, Cell Signaling Technology), mouse anti-ERK1/2 (ab366991, Abcam), rabbit anti-phospho-Akt Ser473 (9271, Cell Signaling Technology), mouse anti-Akt (pan) (2920, Cell Signaling Technology), rabbit anti-EEA1 (3288, Cell Signaling Technology), rabbit anti-Rab7 (9367, Cell Signaling Technology), rabbit anti-Rab11a (2413, Cell Signaling Technology), mouse anti-GAPDH (CB1001, Merck), rabbit phospho-Rb Ser807/811 (8516, Cell Signaling Technology).

Secondary antibodies: IRDye 680 donkey anti-mouse IgG, IRDye 800 donkey anti-rabbit IgG, (LI-COR Biosciences, Lincoln, NE); Alexa Fluor 647 chicken anti-rabbit IgG, Alexa Fluor 647 donkey anti-goat IgG, Alexa Fluor 568 donkey anti-rabbit IgG (Life Technologies, Darmstadt, Germany).

### Plasmids

The constructs, PTPRG-mCitrine/-mTFP/-mCherry, PTPN2-mCitrine/-mTFP and EGFR-mCitrine/-mTFP/-mCherry, PTB-mCherry were generated as described previously (Stanoev *et al*, 2018).

### Cell culture and transfections

MCF7 cells [86012803, ECACC] were cultured in Dulbecco’s modified Eagle’s medium (DMEM; PAN-Biotech GmbH, Aidenbach, Germany) supplemented with 10% fetal bovine serum (FBS; PAN-Biotech GmbH), 2 mM L-Glutamine and 1% nonessential amino acids (NEAA, PAN-Biotech GmbH). Cells were transfected 24 h prior to the experiment with cDNA of interest with Fugene6 following the protocol of the manufacturer (Roche Diagnostics, Burgess Hill, UK). MCF10A cells [CRL-10317, ATCC] were grown in DMEM/F12 (PAN-Biotech GmbH) supplemented with 5% horse serum (Sigma-Aldrich Chemie GmbH, Munich, Germany), 20 ng/ml EGF, 500 ng/ml hydrocortisone (Sigma-Aldrich Chemie GmbH), 100 ng/ml cholera toxin (Sigma-Aldrich Chemie GmbH), 10 μg/ml insulin (Sigma-Aldrich Chemie GmbH). HT29 cells (HTB-38, LGC Genomics GmbH) were cultured in Ham’s F12 culture medium (PAN-Biotech GmbH) supplemented with 2 mM L-Glutamine, 1% (NEAA) and 10% FBS. Cells were transfected with Lipofectamine 3000 following the protocol of the manufacturer (Life Technologies). The culture medium was exchanged 7 h after transfection. All cells were grown at 37°C in 5% CO_2_ and regularly checked for mycoplasma (MycoAlert mycoplasma detection kit, Lonza, Basel, Switzerland). Identity of all cell lines was confirmed by STR analysis (Leibniz Institute DSMZ GmbH, Braunschweig, Germany). Cells were serum starved for at least 6 h prior to the experiment with respective culture media containing 0.5% FBS. The consistency of plasmid expression throughout different experiments was ensured by transfecting 80 ng of each plasmid per well of an 8-well Lab-Tek™ chambered cover glass slides (0.8 cm^2^ surface area, ThermoFisher) and the amount of plasmid was scaled up proportional to the culture-well surface area for other culture-plates used. EGFR and PTPs were co-transfected with 1:1 DNA ratio. In PTB-FLIM experiments, a ratio of 1:2 was used for EGFR-mCitrine:PTB-mCherry.

### Generation of gene knock-outs using CRISPR/CAS9

pSpCas9(BB)-2A-GFP (PX458) was a gift from Feng Zhang (Addgene plasmid #48138; http://n2t.net/addgene:48138; RRID:Addgene_48138). The oligonucleotides containing CRISPR guide RNA sequences (sgRNA) were designed (https://portals.broadinstitute.org/gppx/crispick/public) and synthesized (Sigma-Aldrich Chemie GmbH). sgRNA sequences used to create CRISPR KO cell lines are: PTPRG exon 7: 5’-TCCACTATTTCGCTACACGG-3’; PTPN2 exon 6: 5’-AGGGACTCCAAAATCTGGCC-3’; CYBA exon 2: 5’-GTAGGCACCAAAGTACCACT-3’. The sgRNAs were cloned into the BbsI site of the pX458 expression vector as described previously (Ran *et al*, 2013). MCF7 cells were plated on 6 well tissue culture dishes (SARSTEDT AG & Co. KG, Nümbrecht, Germany) and transfected with 2 μg of each pX458 construct using Fugene6. Cells were FACS sorted 24 h post-transfection using GFP expression and single cells were sparsely seeded on 15 cm dishes to form separated clonal colonies. Single clone derived colonies were picked, expanded and evaluated for knockout by western blot analysis.

### EGF-Alexa647 and EGF-DyLight405

The His-CBD-Intein-(Cys)-hEGF-(Cys) plasmid was kindly provided by Prof. Luc Brunsveld, University of Technology, Eindhoven. Human EGF was purified from E. coli BL21 (DE3) and N-terminally labelled with Alexa647-maleimide or DyLight405-Maleimide as described previously (Sonntag *et al*, 2014) and stored at −20°C in PBS.

### Synthesis and purification of DyTO

DyTo was formulated by labeling DYn2 with atto-590azide by azide-alkyne Huisgen cycloaddition (Rostovtsev *et al*, 2002). The reaction was performed by mixing the following reagents at the mentioned final concentrations to obtain 1 ml reaction volume. atto-590azide (6 mM, ATTO-TEC GmbH, Siegen, Germany) dissolved in DMSO was mixed with aqueous solution of click-chemistry grade CuSO_4_ (40mM, Jena Bioscience GmbH, Germany) and TCEP (40 mM, Sigma-Aldrich Chemie GmbH) for 5’. TBTA (20 mM, Sigma-Aldrich Chemie GmbH) was added to the mixture followed by the addition of DYn2 (18 mM, Cayman Chemical, MI, USA) dissolved in DMSO. The reaction was allowed to proceed overnight at room temperature in dark on a rocking platform.

DyTo was separated from the unreacted reagents and by-products by mass-directed preparative HPLC (infinity prep II, Agilent Series 1260, LC-MSD) using reversed-phase C18 column (VP10/125 5 μm) with a constant flow of 20 ml/min. Water/Acetonitrile (H_2_O+0.1% v/v TFA, CH_3_CN+0.1% v/v TFA) system was used as eluent. The product identity was verified on mass spectrometer integrated with HPLC system. The DyTo fraction obtained from the HPLC was subject to reduced-pressure evaporation and the final product was obtained as dry-powder that was stored at −20°C.

### Immunoprecipitation and western blots

Cells grown in 6-well tissue culture plates (SARSTEDT) were treated as per the experimental requirement and lysed using RIPA cell lysis buffer (50 mM Tris-HCl pH 7.9, 150 mM NaCl,1% IGEPAL, 0.5% Na-deoxycholate, 20 mM NEM) supplemented with Complete Mini EDTA-free protease inhibitor (Roche Applied Science, Heidelberg, Germany) and phosphatase inhibitor cocktail 2 and 3 (1:100, P5726 and P0044, Sigma-Aldrich). For immunoprecipitation (IP), equal amounts of protein lysates were incubated with pull-down antibody overnight at 4°C followed by 2 h treatment with Dynabeads Protein G magnetic beads (10004D, ThermoFisher, MA) for pull down. Total and IP protein were prepared in XT Sample Buffer (1610791, Bio-Rad, CA) supplemented with 0.05 M DTT. The protein bands were resolved by SDS/PAGE using NuPAGE Novex 4-12% Bis-Tris gels (ThermoFisher) in MOPS-SDS running buffer (ThermoFisher) at 200V constant voltage, blotted to polyvinylidene difluoride membrane by wet-tank transfer for 1.5 h at 100V constant voltage and blocked with intercept (TBS) blocking buffer (LI-COR Biosciences) for 1 h.

Primary antibody incubation was performed over night at 4°C, followed by washing with TBS/T and incubation with the respective secondary antibodies for 1 h in the dark. After final washing with TBS/T, the blot was scanned using an Odyssey Infrared Imaging System (LI-COR). The integrated intensities of the protein bands on western blots were measured using Fiji (Schindelin *et al*, 2012) and signals were normalized either as the ratio of phosphorylated/oxidized protein to total protein or in case of co-IP 5 experiments as co-IP proteins to corresponding IP proteins.

### Confocal laser scanning microscopy (CLSM)

Cells were cultured and transfected (as described above) for CLSM experiments on 4 or 8-well Lab-Tek™ chambered cover glass slides (ThermoFisher). Before imaging, the culture media was replaced with phenol red-free DMEM supplemented with 25 mM HEPES (PAN-Biotech GmbH). Confocal images were acquired using a Leica TSC SP8 microscope (Leica Microsystems, Wetzlar, Germany), equipped with an environment-controlled chamber (Life Imaging Services) set to 37°C, a 405-nm diode laser and a white light laser (white light laser Kit WLL2, NKT Photonics). Imaging was done with HC PL APO CS2 63x/1.4NA oil immersion objective and pinhole between 1 to 1.7 airy units. Following excitation wavelengths were used for proteins with fluorescent fusion tags/labels: BFP/DyLight 405 (405 nm), TFP (458 nm), mCitrine (514 nm), mCherry/Alexa Fluor 568 (561 nm), Atto-Tec 590 (593 nm), Alexa Fluor 647 (640 nm). Fluorescence emission was detected by hybrid detectors (HyD) restricted at: BFP/DyLight 405 (415-450 nm), TFP (468-485 nm), mCitrine (524-560 nm), mCherry/Alexa Fluor 568 (571-620 nm), Atto-Tec 590 (605-630 nm), Alexa Fluor 647 (655-690 nm). 12-bit images were recorded by frame-by-frame sequential scanning at 512×512 pixels, with 80 Hz scanning frequency.

### Immunofluorescence

Cells grown in 4 or 8-well Lab-Tek™ chambered cover glass slides (ThermoFisher) were fixed with 4% PFA (15710, EM-grade, EMS, Hatfield, PA) in TBS (v/v) for 10’. Cells were permeabilized with 0.2% Triton X-100 (v/v) for 8’. Background staining was blocked by 1 h incubation with Intercept (TBS) Blocking Buffer at room temperature. Samples were incubated with primary antibodies at 4°C overnight and secondary antibodies for 1 h at room temperature. Antibodies were diluted in Intercept (TBS) Blocking Buffer. All washing steps following fixation, permeabilization, primary or secondary antibody incubation were performed with TBS.

The extent of co-localization in CLSM images of the endosomal marker proteins (EMP): Rab11a, EEA1 or Rab7 with the protein of interest (POI): RPTPγ-mCitrine or EGFR-mCherry was computed in Fiji (Schindelin *et al*, 2012) by defining ROIs in EMP and POI channels by intensity thresholding in background-subtracted images. The EMP mask was converted to a binary mask and multiplied with the thresholded POI image. The integrated intensity from this overlapping area was normalized to the total integrated intensity of POI in the whole cell to obtain the fraction of POI co-localized to the EMP.

### Fluorescence lifetime imaging microscopy (FLIM)

FLIM of EGFR-mCitrine/PTB-mCherry or PTP-mCitrine/DyTo was performed at 37°C on the Leica SP8 laser-scanning microscope equipped with a fast lifetime contrast module (FALCON, Leica Microsystems) using the 63×/1.4NA oil objective. mCitrine was excited using the white light laser at a frequency of 20 MHz and wavelength of 514 nm, and fluorescence emission was collected between 525 to 560 nm on hybrid detectors (HyDs). The photon collection was split amongst 2-3 HyDs with an AcoustoOptical Beam Splitter (AOBS). Photons were integrated for a total of approximately 15-20 s per image (~300-600 photons/pixel, sum of all detectors) using the FALCON system.

The donor count images were thresholded above the background fluorescence. Global analysis of FLIM-FRET images was performed as described previously (Grecco *et al*, 2009) by a custom script implemented in Python to deduce the spatial distribution of the fraction of FRET-exhibiting donor (α) for EGFR-phosphorylation (PTB-bound fraction of EGFR-mCitrine: ap) and PTP-oxidation (DyTo-bound fraction of PTP-mCitrine: α_ox_).

### Fluorescence photoactivation microscopy

Fluorescence photoactivation microscopy was carried out at 37°C on a Leica SP8 microscope using a 63×/1.4NA oil objective. Cells were transfected with photoactivatable paGFP-RPTPγ and RPTPγ-mCherry, with or without co-expression of BFP-Rab11a. Pre-and post-photoactivation images were acquired for paGFP-RPTPγ and RPTPγ-mCherry using 488 nm and 561 nm laser excitation, respectively. For photoactivation, paGFP was excited with the 405 nm diode laser at 80% transmission in 3 frames on perinuclear region of interest (ROI) corresponding to the RE, identified by localization of RPTPγ-mCherry. In the post-activation step, fluorescence images for paGFP-RPTPγ and RPTPγ-mCherry were acquired with a time interval of 30”, using the 488 nm and 561 nm WLL laser at 10% power. At the end of the experiment an image of BFP-Rab11a was acquired with 5% 405-laser power. Following background correction, fluorescence gain on the PM was quantified by the ratio of local (cell periphery) paGFP-RPTPγ to RPTPγ-mCherry fluorescence using Fiji (Schindelin *et al*, 2012).

### Spatial-Temporal Maps

CSLM images obtained at 1’ interval were background corrected prior to further processing. PM and nuclei in individual cells were masked using RPTPγ-mCitrine and CBL-BFP fluorescence, respectively. For each pixel in the cell space, the normalized radial localization (r) was calculated as follows:

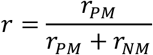

where *r_PM_* and *r_NM_* are the shortest Euclidean distances from each pixel to the PM and NM, respectively. The pixels were segmented in 10 radial bins based on their normalized distances. The mean fluorescence intensities were calculated in each segment for RPTPγ-mCitrine, EGFR-mCherry and EGF-Alexa647 to obtain their spatial distribution profiles at each time point for individual cells. The spatial profiles obtained at consecutive time-points were concatenated to form 3D spatiotemporal maps (STMs) and then combined to yield an average STM (Stanoev *et al*, 2018).

### Fluorescence anisotropy Microscopy

Fluorescence anisotropy microscopy was performed before and during EGF stimulation (160 ng/mL) in WT or RPTPγ-KO MCF7 cells ectopically expressing EGFR-QG-mCitrine (Baumdick *et al*, 2015). Cells were incubated in cell culture medium with 0.5% FCS for at least 6 h before the experiment. The microscope consisted of an Olympus IX-81 body equipped with a MT20 illumination system, an 20x/0.7NA objective and CellR software (Olympus GmbH, Hamburg, Germany). A linear dichroic polarizer (Meadowlark Optics, Frederick, Frederick, CO, USA) was placed in the illumination path of the microscope, and two identical polarizers in an external filter wheel were used to measure parallel (*I*_II_) and perpendicular (*I*_⊥_) polarized emitted light in two separate images. Steady state anisotropy (*r_i_*) was calculated in each pixel *i*:

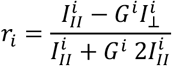

The *G* factor (*G*^i^) was determined by acquiring the ratio of the intensities at perpendicular and parallel orientations for fluorescein in solution, which had a steady-state anisotropy close to zero due to its fast rotation. Quantification of anisotropy was performed using Fiji (Schindelin *et al*, 2012).

### Super-Resolution radial fluctuation (SRRF) under ultrarapid cryo-arrest

Ultra-rapid cryo-arrest of a stable MCF7 cell line expressing SNAP-EGFR and RPTPγ-mCitrine was done as recently described (Huebinger *et al*, 2021). For this, the cells were grown on fibronectin-coated (5 μg/cm^2^, F0895, Sigma-Aldrich) circular microscopy cover slides (No.1; Ø 4 mm) that were mounted to chambers from biocompatible silicone (4-well micro-Inserts; ibidi GmbH, Gräfelfing, Germany). Before the experiment, Snap-EGFR was labeled with 0.5 μM Snap-Surface Alexa647 (New England Biolabs GmbH, Frankfurt, Germany) for at least 60’. The cover slides were mounted to the ultra-rapid cryo-arrest device (Huebinger *et al*, 2021), which was placed on top of an epifluorescence microscope. The cells were cryo-arrested during observation on the microscope. The microscope consisted of an IX-83 microscope body equipped with a 40x/0.95NA objective (UPlanApo) and a MT20 illumination system (Olympus Deutschland GmbH, Hamburg, Germany) as well as an Orca-R2 camera (Hamamatsu Photonics, Hamamatsu, Japan). Series of 100 widefield fluorescence images were acquired for SRRF reconstructions of mCitrine (excitation: 470/40 nm; dichroic mirror 495 nm; emission 520/35 nm) and Alexa647 (excitation: 620/60 nm; dichroic mirror 640 nm; emission 700/75 nm) with pixel length of 163 nm and a frame rate of 1 frame/s. SRRF reconstruction was done using the NanoJ plugin (Laine *et al*, 2019) to Fiji (Schindelin *et al*, 2012).

### Quantification of absolute EGFR-expression

EGF-Alexa647 binding and ectopic EGFR-mCitrine expression was linearly correlated for CLSM image data (Fig EV1D). Absolute EGFR abundance in MCF7 and MCF10A cells is known from the literature (Charafe-Jauffret *et al*, 2006). By application of a 2-point linear calibration curve between median EGF-Alexa647 signal in MCF10A (10^5^ receptors) and Y-intercept corresponding to endogenous EGFR expression in MCF7 wildtype cells (10^3^ receptors), the EGF-Alexa647 signal and corresponding EGFR-mCitrine signals were converted to absolute receptor numbers. From that distribution (Fig EV1D), the absolute number of EGFR/cell was calculated based on measured EGFR-mCitrine intensity in MCF7 WT, RPTPγ-KO and p22^phox^-KO cells at initial timepoints of time series. Due to inter-experimentally changing imaging settings, a conserved median for EGFR/cell in EGFR-mCitrine transfected MCF7 WT cells was assumed and by this a normalization factor was calculated and applied.

### Single cell morphodynamics

WT, RPTPγ-KO and p22^phox^-KO MCF7 cells were seeded on Lab-Tek™ chambered cover glass slides (ThermoFisher), transfected with EGFR-mCitrine or BFP-tkRas and incubated in cell culture medium with 0.5% FBS for at least 6 h before the experiment. Cells were stimulated with sustained doses of EGF-Alexa647 (1 ng/ml or 160 ng/ml) and CLSM images were obtained every 2’ for 60’.

For the analysis of single cell morphodynamics, cells within dense colonies were excluded. Temporal-stacks from EGFR-mCitrine or BFP-tkRas fluorescence images were segmented to binary masks: If necessary, a local contrast normalization (40 px, Integral Image Filters plugin (Schindelin *et al*, 2012)) was performed, and an Otsu automated intensity threshold applied. Masks were manually checked and corrected, cleaned and split into single cell masks, from which area and Perimeter were measured. Morphometric changes were calculated using a Perimeter-Ratio:

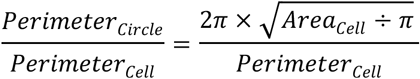

Shape change of cells over time is displayed using the temporal color coder plugin (developed by Kota Miura, Centre for Molecular and Cellular Imaging, EMBL Heidelberg, Germany) on the bare outlines of cell masks.

### Wound healing assay

WT, RPTPγ-KO and p22^phox^-KO MCF7cells were seeded onto fibronectin-coated (1.85 μg/cm^2^) 24-well culture plates containing two-well silicone culture inserts (IBIDI) to create a cell-free gap. After the culture insert was removed, cells were stimulated with EGF-Alexa647 (1 or 160 ng/ml) and imaged immediately. Transmission images were acquired on the Leica SP8 microscope with the environment-controlled chamber maintained at 37°C and 5% CO_2_ using 10×/0.3 or 0.4NA air objective for 12 h at a time interval of 10’.

To analyze cell-migration, segmentation was performed using Trainable Weka Segmentation Plugin in Fiji (Arganda-Carreras *et al*, 2017). For each experiment, one image from the transmission stack was chosen to manually label the pixels in 3 classes: cell-free gap, inner part of the cells and cell boundaries. Considering the best trade-off between precision and recall for a single class, Bayes Net Classifier was trained on the labelled data and then applied to the whole image stack to obtain masks for each class for the time series. The cell-count was quantified with MATLAB/Python by size thresholding the detected particles for noise removal, finding geometric centers of the objects, dividing every 512×512 image on 16×16 bins of size 32×32 pixels and calculating the number of cells in the constructed bins. The number of cells in each bin was averaged for 1 h. The boundary bin position was detected via the largest drop of cell density on the first frame. The bins in the gap and in the bulk were defined with respect to the position of the gap boundary (Fig 5G).

### Clonogenic assay

WT, RPTPγ-KO and p22^phox^-KO MCF7 cells were uniformly seeded in linear increment (100, 200 or 300 cells per well) in 6-well tissue culture plates (SARSTEDT), incubated at 37°C, 5% CO_2_ in DMEM growth media supplemented with 10% FBS, 2 mM L-Glutamine and 1% NEAA with media exchange every alternate day. After 7 days, the plates were washed with PBS, fixed with 4% (v/v in PBS) PFA (EM-grade, EMS) for 10’ and stained with 0.05% (v/v in PBS) crystal violet (Sigma-Aldrich Biochemie GmbH) for 10’. The excess crystal violet was removed by washing three times with PBS. The lids were removed for 3-4 h to dry the plates before imaging. Plates were scanned on Typhoon TRIO+ variable mode imager (GE Healthcare, Buckinghamshire, UK) with the following settings: Cy5 filter (ex. 633nm, em. 670 band-pass and PMT 530), pixel size 50 μm and focal plane +3 mm.

To quantify the occupied culture-well area by proliferated cell-colonies, the image of the scanned well was intensity thresholded for the pixels occupied by colonies and a binary mask was obtained. By using Analyze Particles command in Fiji (Schindelin *et al*, 2012), the area of the mask was calculated and normalized to the area of the culture well.

### Computational modelling and data fitting

To capture the interactions between EGFR and the PTPs across the different conditions, the following model of differential equations was used:

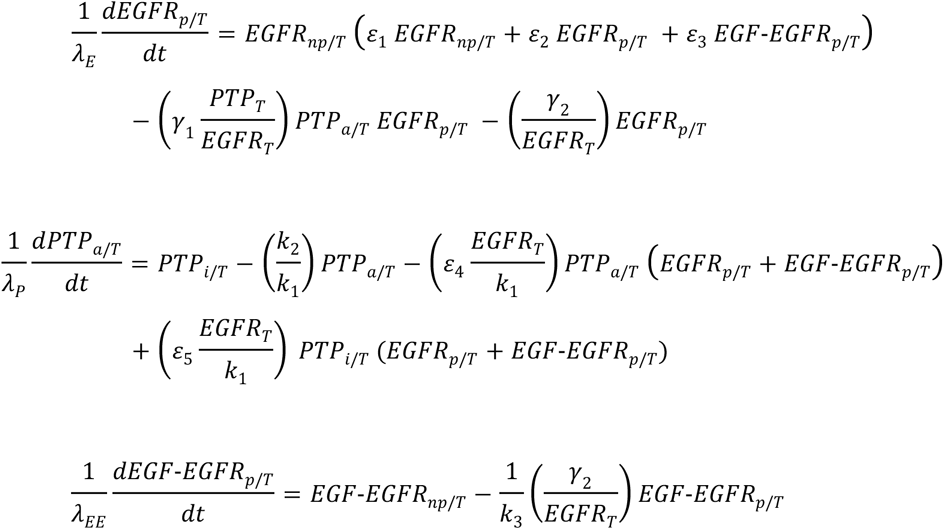

Here, *EGFR_p/T_, PTP_a/T_* and *EGF-EGFR_p/T_* describe the fractions of phosphorylated/active proteins, relative to the respective total protein concentration, while 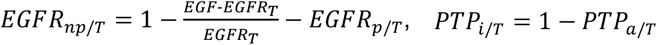 and 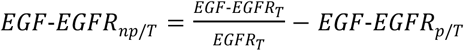 describe the fractions of non-phosphorylated/inactive proteins. Parameters *ε*_1_, *ε*_2_ and *ε*_3_ represent the second-order phosphorylation rate constants of the unliganded EGFR, while *k*_3_ represents the first-order phosphorylation rate constant of the liganded EGFR. These four EGFR activation parameters are assumed to be dependent on the kinetic constants of EGFR only, therefore their values are shared across the different conditions, while the remaining parameters on the right side are fitted in condition-specific manner. The rest of the rate constants depict: *γ*_1_-second-order PTP-specific dephosphorylation, *γ*_2_-first-order generic PTP dephosphorylation, *k*_2_/*k*_1_-intrinsic PTP deactivation/activation ratio, *ε*_4_ and *ε*_5_ – second-order EGFR-dependent deactivation and activation respectively, while *EGFR_T_* and *PTP_T_* depict the respective protein concentrations. The bracketed parameter groups were treated as single parameters. For a given parameter set, bifurcation profiles depicting the dependence of the fraction of phosphorylated EGFR *EGF-EGFRT_p/T_* + *EGF-EGFR_p/T_*) on the input fraction of EGF-bound 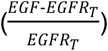 in steady-state were calculated, using a custom-made continuation algorithm. The resulting profiles were compared (using sum-of-squares) with the corresponding dose-response dependencies from the experimental data points (Fig EV1K; offsets in data were removed except for the conditions with pre-activation). Finally, a custom-made Metropolis Hastings algorithm was used to perform parameter estimation of all the parameters together. Bifurcation diagrams were then generated depicting the dependence of the fraction of phosphorylated EGFR (*EGFR_p/T_* + *EGF-EGFR_p/T_*) (α_p_) on the PTP-dependent dephosphorylation rate 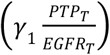 in absence of EGF-bound EGFR, marking the estimated parameter value (Fig EV1L). These bifurcation profiles, calculated additionally for increasing input fractions of EGF-bound EGFR 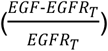, were used to compile the 3D bifurcation diagrams (Fig 1F-H). The estimated parameters for the different conditions are shown in Table 1.

**Table 1.**
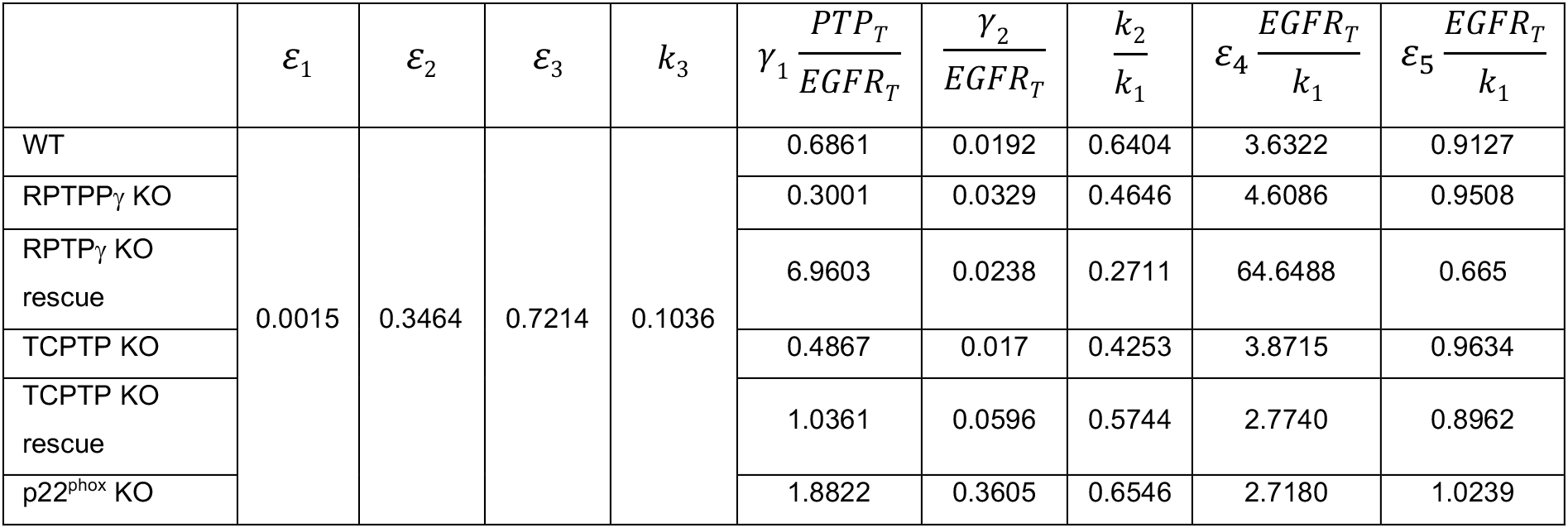
Estimated parameters from bifurcation analysis of indicated cell lines

## Acknowledgments

We would like to thank Dr. A. Krämer for critically reading the manuscript, M. Reichl for cell culture work and Dr. S. Müller for help with microscopy and Dr. M. Schulz for FACS (all at Systemic Cell Biology, MPI for Molecular Physiology). We thank Dr C. Sorio, Department of Pathology, University of Verona, Verona, Italy for the generous gift of rabbit anti-RPTPγ.

## Author contributions

PIHB conceived the project and wrote the manuscript with MSJ and help from JH. MSJ performed all experiments except: FLAP and corresponding analysis by AS, Cryo-nanoscopy and fluorescence anisotropy by JH. AS performed bifurcation analysis based on concepts of PIHB. BS analyzed morphodynamics and VZ wound healing assays. LR and MSJ generated CRISPR-Cas9 KO cell lines. KM validated KO cell lines and performed EGF conjugation reactions.

## Data and materials availability

All data available in the main text or supplementary materials. Correspondence and request for materials should be addressed to Philippe I. H. Bastiaens. pSpCas9(BB)-2A-GFP (PX458) (Addgene plasmid # 48138) plasmids require a material transfer agreement from Addgene.

