## Supplemental information for "RPTPγ is a redox-regulated suppressor of promigratory EGFR signaling"

Expanded view material for

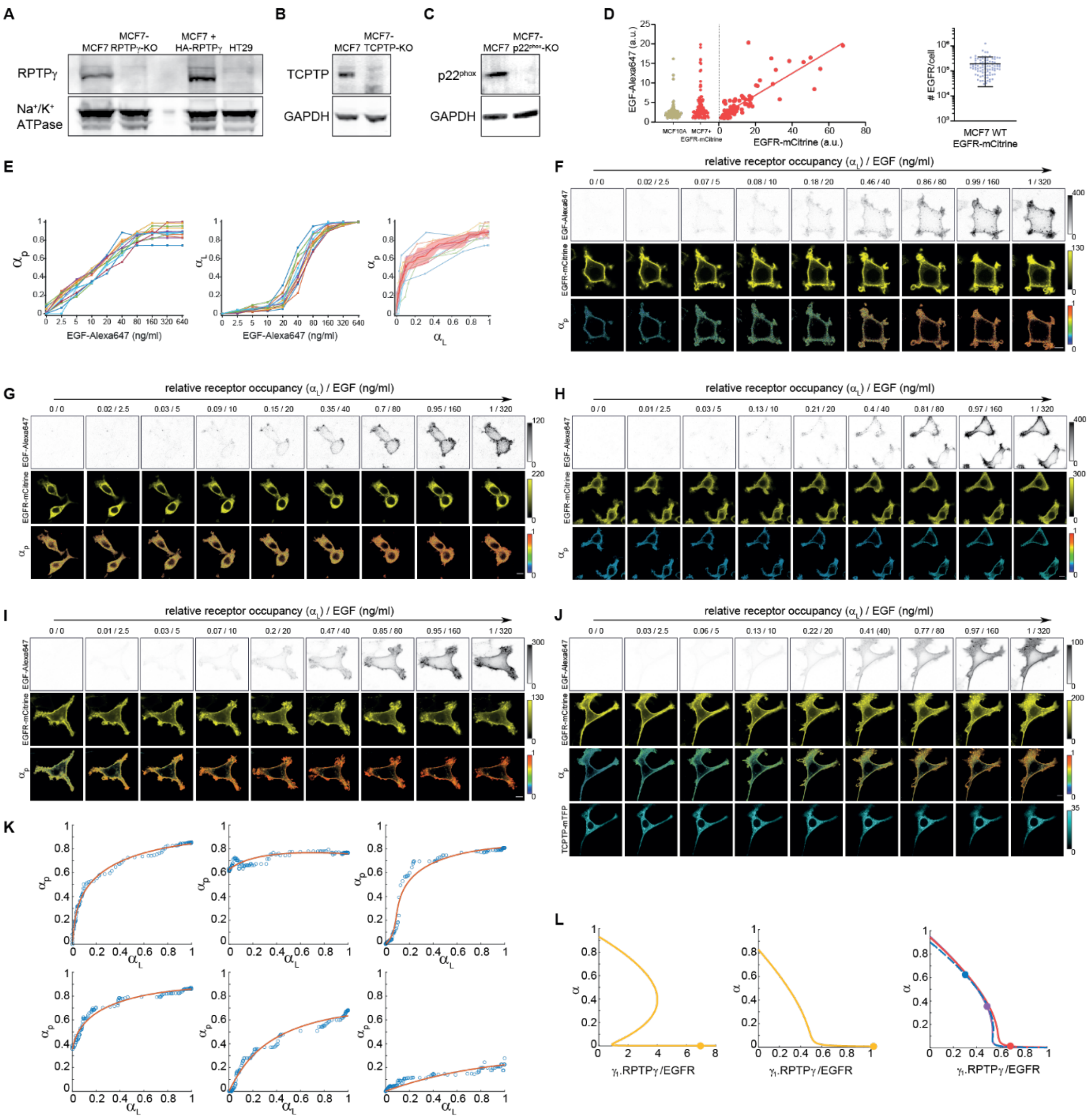


**Figure EV1. Perturbation of major PTP activities impacts the growth factor response of EGFR**

A. Western blot against endogenous or exogenous (HA-RPTPγ) RPTPγ (top row) in cell lysates obtained from membrane protein extraction of MCF7 cells without (first lane) or with (second lane) CRISPR-Cas9-mediated RPTPγ-KO and from RPTPγ deficient HT29 cells (right lane). Bottom row: $Na^{+}/K^{+}$ ATPase loading control

B, C. Knockout of TCPTP and $p22^{phox}$ respectively (top rows) with GAPDH as a loading control (bottom rows).

D. Left panel: comparison of EGF-Alexa647 binding in endogenous EGFR expressing MCF10A (beige) to exogenous EGFR-mCitrine expressing MCF7 cells (red), along with the levels of EGF-Alexa647 binding as a function of EGFR-mCitrine expression in MCF7 cells, right panel: Derived number of EGFR per cell in MCF7 WT expressing ectopic EGFR-mCitrine.

E. Response of individual EGFR-mCitrine expressing MCF7 cells to cumulative doses of EGF-Alexa647 (2.5-640 ng/ml). Left: fraction of phosphorylated EGFR-mCitrine measured by FLIM ($\alpha_p$) as a function of administered EGF-Alexa647 dose, middle: fraction of EGF-Alexa647 binding to EGFR-mCitrine (receptor occupancy $\alpha_L$) upon each administered dose, right: $\alpha_p$ plotted against $\alpha_L$. Colored thin lines: individual cells; Solid red line with shaded bounds: Moving medians with median absolute.

F. Representative *in cell* dose response fluorescence micrographs after cumulative EGF-Alexa647 administration (0–320 ng/mL) at 1.5' interval. First row: EGF-Alex647; Second row: EGFR-mCitrine; Third row: phosphorylated EGFR fraction ($\alpha_p$) in MCF7 WT cells.

G. Same as (F) for RPTP$\gamma$-KO.

H. Same as (F) for p22$^{phox}$-KO.

I. Same as (F) for TCPTP-KO.

J. Same as (F) for TCPTP-KO with TCPTP-mTFP (fourth row) ectopic expression.

Data information: In (F-J), all scale bars: 10 $\mu$m. N=3-4, n=12-14.

K. Dose-response models fitted to the experimental data depicting the dependence of the fraction of phosphorylated EGFR ($\alpha_p$) on the fraction of EGF-bound EGFR ($\alpha_L$). Upper row: MCF7 WT (left), RPTP$\gamma$-KO (middle), RPTP$\gamma$-KO with RPTP$\gamma$-mTFP expression (right) and Lower row: TCPTP-KO (left), TCPTP-KO with TCPTP-mTFP expression (middle), p22$^{phox}$-KO (right).

L. 2D-bifurcation diagrams showing the dependence of EGFR phosphorylation ($\alpha_p$) on RPTP$\gamma$/EGFR expression ratio. Left: RPTP$\gamma$-KO with RPTP$\gamma$-mTFP expression, middle: TCPTP-KO with TCPTP-mTFP expression, right: WT (red), RPTP$\gamma$-KO (blue dotted) and TCPTP-KO (purple) MCF7 cells. Solid circles: Poising of the system prior to EGF-stimulus.

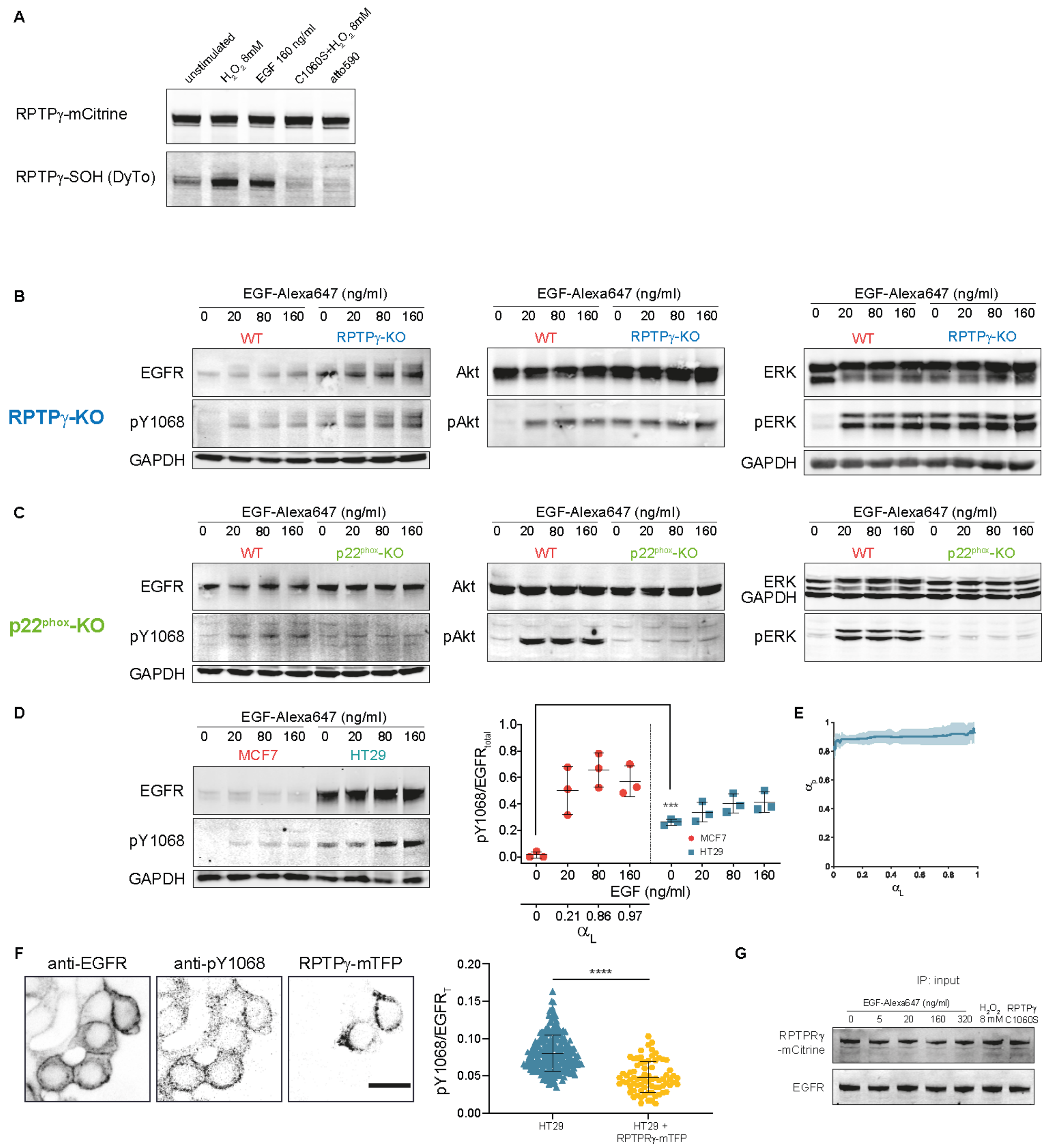


**Figure EV2. Redox regulated RPTPγ prevents aberrant signaling of EGFR**

A. Representative IP-western blot showing co-IP of oxidized RPTPγ bound to DyTo (bottom row) upon RPTPγ-mCitrine (upper blot: lanes 1-3, 5) or RPTP$\gamma^{C1060S}$-mCitrine (lane 4) pull-down by anti-GFP antibody from cell lysates of MCF7 cells that were unstimulated (lane 1) or upon subject to 10' stimulation with $H_2O_2$ (8 mM; lane 2) or EGF-Alexa647 (160 ng/ml; lane 3). Lane 5: Lysates treated with bare Atto590-azide (without DYn2-coupling).

B. Representative western blots showing EGFR (left), Akt (middle) and Erk (right) in the top rows with corresponding phosphorylation (middle row) in WT and RPTP$\gamma$-KO MCF7 cells, prior to and upon 5' stimulation with EGF-Alexa647 (20, 80 and 160ng/ml). GAPDH (bottom row) was used as a loading control.

C. Same arrangement as (B) for WT and p22$^{phox}$-KO MCF7 cells.

D. Left: Representative western blot showing phosphorylated EGFR at tyrosine 1068 (pY1068) in MCF7 WT and HT29 cells, prior to and upon 5' stimulation with EGF-Alexa647 (20, 80 and 160ng/ml). Right: Quantification with mean±SD, N=3, ***P<0.001: unpaired two-tailed t-test.

E. Phosphorylation response of EGFR-mCitrine to cumulative EGF-Alexa647 doses (0-320 ng/ml) in HT29 cells (N=3, n=12). Solid line: moving median from single cell profiles; shaded bounds: median absolute deviations.

F. Confocal micrographs (left panel) and quantification (right panel) of immunostained endogenous EGFR (left image), phosphorylated EGFR at tyrosine 1068 (middle image: pY1068) and RPTP$\gamma$-mTFP fluorescence (right image) in HT29 cells without EGF-stimulus. Scale bar: 10 $\mu$m. N=3, n>75, ****p<0.0001: unpaired two-tailed t-test.

G. Loading control western blot for the IP in Fig 3D.

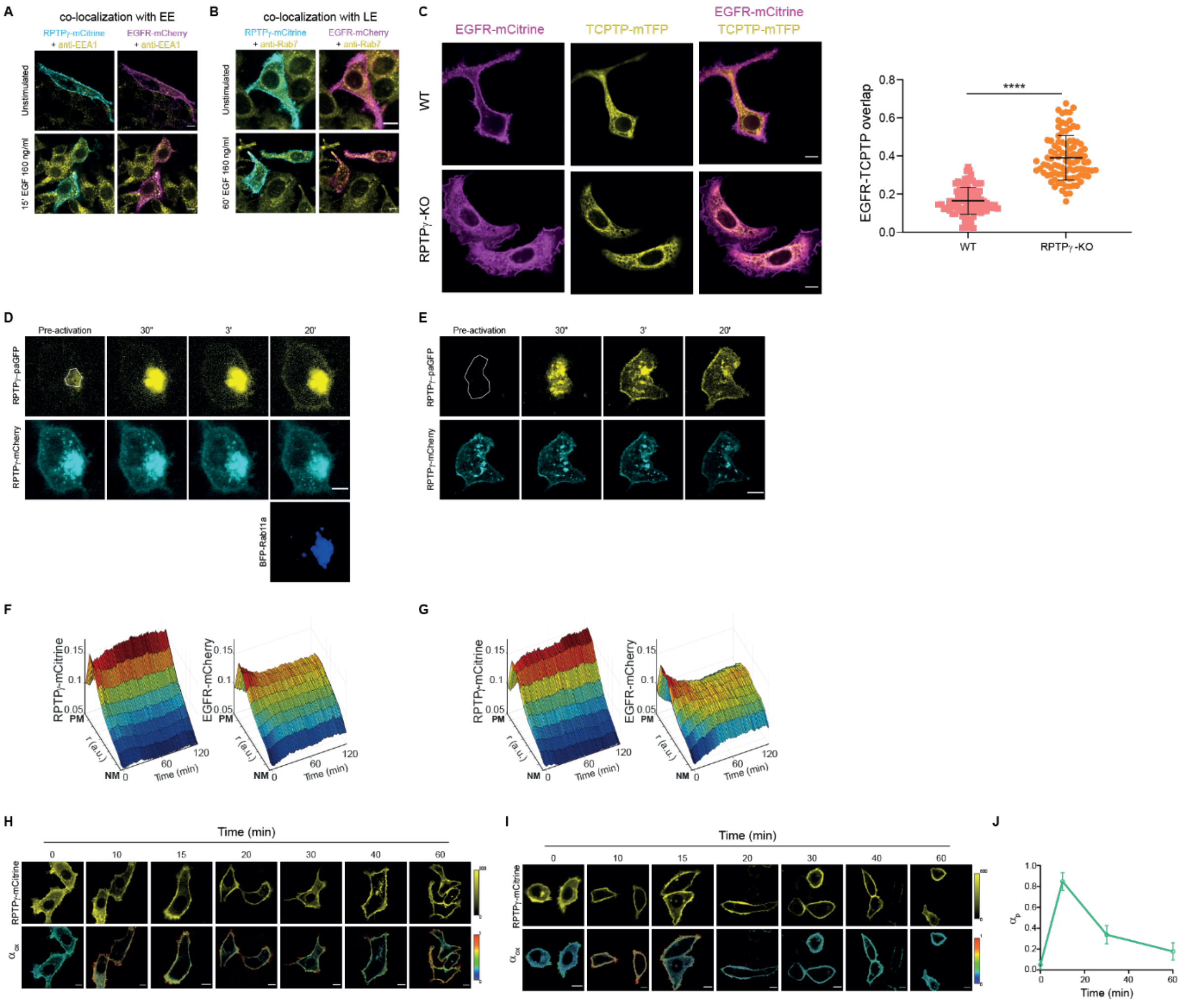


**Figure EV3. Recycling of EGFR-RPTPγ regulates RPTPγ oxidation**

A. Confocal micrographs of MCF7 WT cells showing the co-localization of RPTPγ-mCitrine (cyan, first column) and EGFR-mCherry (magenta, second column) with early-endosome marked by immunostaining against EEA1 (yellow), prior to (top row) or after EGF-DyLight405 stimulus (160 ng/ml, bottom row).

B. Same as (A) with late-endosome marked by immunostaining against Rab7 (yellow).

C. Left panel: Representative confocal micrographs comparing the steady state co-localization of EGFR-mCitrine (magenta) and the ER-marker TCPTP-mTFP (yellow) in WT (top row) to RPTPγ-KO (bottom row) MCF7 cells. Right panel: Quantification of the fraction of EGFR-mCitrine co-localizing with ER-marker. N=3, n>50. ****P<0.0001: unpaired two-tailed t-test.

D. Representative confocal micrographs for the fluorescence photoactivation of paGFP-RPTPγ (top row) on the recycling endosome (white-rimmed region) in MCF7 cells with co-expressed RPTPγ-mCherry (middle row) and BFP-Rab11a (last row) at indicated times after photoactivation. Gamma correction for all channels: 0.18.

E. Same as (D) without BFP-Rab11a co-expression.

F. Average spatial-temporal maps constructed from confocal micrographs obtained at 1' interval from live MCF7 cells showing the distributions of RPTPγ-mCitrine and EGFR-mCherry as a function of their radial distance and time, upon sustained low (20 ng/ml) EGF-Alexa647 stimulus (N=3, n=13 cells)

G. Same as (F) with sustained high (160 ng/ml) EGF-Alexa647 stimulus (N=3, n=14).

H. Representative confocal micrographs of MCF7 cells expressing RPTPγ-mCitrine (top row) and its oxidized fraction estimated using DyTo-FLIM ($\alpha_{ox}$, bottom row), upon sub-saturating (20 ng/ml) sustained EGF-Alexa647 stimulus at the indicated time points.

I. Same as (H) for saturating (160ng/ml) sustained EGF-Alexa647 stimulus.

J. Temporal profile of EGFR-mCitrine phosphorylation ($\alpha_p$) in MCF7 WT cells post 5' pulsed stimulus with saturating EGF-Alexa647 (160 ng/ml). mean±SD, N=3, n=24. All scale bars: 10 μm.

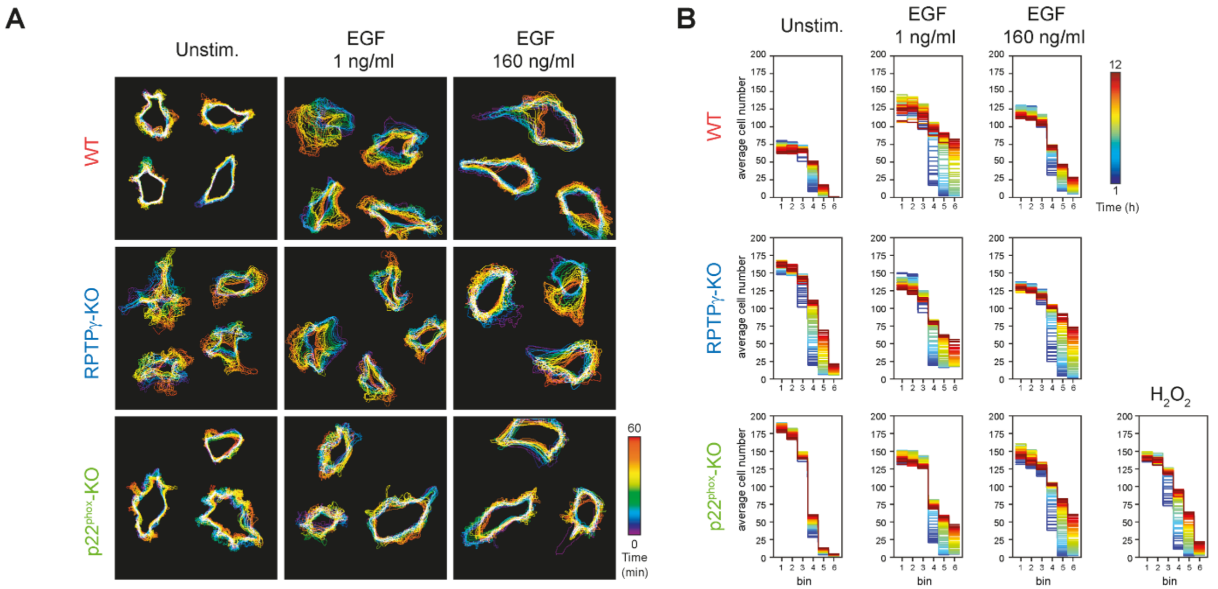


**Figure EV4. RPTPγ suppresses EGFR PM-activity associated migration but not endosomal-activity induced proliferation**

A. Representative cell contour maps showing the temporal changes (color code: lower right) in the cell morphology for WT (upper row), RPTPγ-KO (middle row) and p22$^{phox}$-KO (bottom row) MCF7 cells expressing EGFR-mCitrine without (first column) or during EGF-Alexa647 stimulus (1 ng/ml, second column; 160 ng/ml, third column) over 60'.

B. Temporal color-coded (time (h): color bar upper right) maps depicting the dynamics of migrating cells across the spatial bins upon removal of the migration-barrier. Average number of cells at each hour is plotted for 6 spatial bins under consideration (location of the bins in the migration chamber depicted in Fig 5G: right panel). N=4-5 per condition.

**Movie EV1. Time-lapse wound healing assay**

Time-lapse transmission microscopy of WT (first row), RPTP$\gamma$-KO (second row), p22$^{phox}$-KO (third row) MCF7 cell invasion into a cell-free gap after removal of migration barrier for different stimuli conditions as indicated.
